# Genome-wide synthetic-lethality screen of Bam complex associated genes in *Escherichia coli*

**DOI:** 10.1101/2023.07.05.547807

**Authors:** Jack A. Bryant, Kara A. Staunton, Hannah M. Doherty, Micheal B. Alao, Xuyu Ma, Joanna Morcinek-Orłowska, Emily C. A. Goodall, Jessica Gray, Mathew Milner, Luke Kidger, Charly D. Neilson, Jeffrey A. Cole, Felicity de Cogan, Timothy J. Knowles, Monika Glinkowska, Danesh Moradigaravand, Ian R. Henderson, Manuel Banzhaf

## Abstract

Biogenesis of the bacterial outer membrane is key to bacterial survival and antibiotic resistance. Central to this is the β-barrel assembly machine (Bam) complex and its associated chaperones, which are responsible for transport, folding and insertion of outer membrane proteins (OMPs). The *Escherichia coli* Bam complex is composed of two essential subunits, BamA and BamD, and three non-essential accessory lipoproteins, BamB, BamC and BamE. Optimal Bam function is further dependent on the non-essential periplasmic chaperones DegP, Skp and SurA. Despite intensive study, the specific function of these non-essential Bam-associated proteins is not fully understood. Here, we analysed Δ*bamB*, Δ*bamC*, Δ*bamE*, Δ*surA*, Δ*skp* and Δ*degP* knockout strains by phenotypic screening, conservation analysis and high-throughput genetics. We identify hundreds of synthetic-lethal interactions and reveal that Bam complex activity is impacted by changes in outer membrane lipid composition and that enterobacterial common antigen is essential in the absence of the chaperone SurA. We also show genes responsible for synthesis of peptidoglycan are synthetically-lethal with Bam accessory lipoprotein encoding genes. Together, our data indicates potential mechanisms for coordination of OMP biogenesis with other cellular growth processes such as LPS and peptidoglycan biogenesis.

## Introduction

The Gram-negative outer membrane is instrumental for virulence, antimicrobial resistance, and immune evasion. The outer membrane is an asymmetric lipid bilayer consisting of phospholipid on the inner leaflet, lipopolysaccharide (LPS) on the outer leaflet, integral outer membrane proteins (OMPs) and associated lipoproteins. Biogenesis of the outer membrane is completed by several multi-component proteinaceous machines. Central to this is the β-barrel assembly machine (Bam) complex, the essential protein machinery responsible for folding and insertion of OMPs into the outer membrane, which then act as transporters, porins, receptors, virulence factors, adhesins and enzymes [1, 2]. All OMPs are synthesised in the cytoplasm before being trafficked through the inner membrane to the periplasmic space by the Sec machinery. OMPs are prone to aggregation and must be maintained in an unfolded state during their translocation to the Bam complex [3]. Three quality control proteins that chaperone OMPs across the periplasm are SurA, Skp and DegP, the latter of which also serves as a protease to degrade misfolded OMPs in the periplasm. The double mutants Δ*surA*Δ*skp* and Δ*surA*Δ*degP* are not viable, which suggests that Skp, SurA and DegP are functionally redundant [4]. The primary chaperone pathway to the Bam complex is thought to be SurA, while Skp and DegP are suggested to form a secondary pathway that is amplified in stress conditions [5]. However, single molecule studies with purified OmpC and Skp or SurA demonstrate that the two proteins recognise different conformations of the OMP and that while Skp is capable of dispersing aggregated OmpC, SurA is not [6]. In addition, the absence of SurA increases degradation of OMPs that have stalled on the Bam complex, whereas loss of Skp in this scenario has the opposite effect, enhancing their assembly [7]. This is potentially due to the role of Skp in recognising and removing stalled OMP substrates from the Bam complex before being degraded alongside the misfolded OMP by the protease DegP [8]. These studies suggest that Skp and SurA have specialised roles in maintaining OMPs in a folding competent state during periplasmic transit.

Following transport across the periplasmic space, OMPs are received at the outer membrane by the Bam complex. In *Escherichia coli*, the Bam complex is composed of two essential subunits, the outer membrane β-barrel BamA and the lipoprotein BamD, and three non-essential accessory lipoproteins, BamB, BamC and BamE [9, 10]. BamA is the central component of the complex and consists of a 16-stranded β-barrel and five periplasmically localised POlypeptide-TRansport-Associated (POTRA) domains that function to dock the accessory subunits [11, 12]. The POTRA domains are hypothesised to interact with incoming unfolded OMP substrates and feed them into the assembly machinery [13]. BamD is a lipoprotein composed of five tetratricopeptide repeat (TPR) domains that interact with POTRA 5 of BamA [9, 14, 15]. The remaining components of the complex are the accessory lipoproteins, BamB, BamC and BamE, which interact with BamA and BamD to form a functionally coordinated ring structure under the BamA barrel on the periplasmic face of the outer membrane [16–20].

The Bam complex accessory lipoproteins each contribute to full activity of the complex through a shared redundant function of coordinating efficient BamA/D interaction in OMP engagement and folding. Beyond this shared function, there is some limited information about specific roles of these proteins in the cell [10, 21]. For example, as part of the Rcs stress response signalling pathway, the outer membrane lipoprotein RcsF is threaded through an OMP during folding by the Bam complex. This process specifically requires BamE to directly interact with and coordinate BamA and BamD to enable completion of OMP folding around RcsF [22–24]. Lethal jamming of the Bam complex in the absence of BamE can be prevented by BamB [23, 25]. BamB has also been shown to be important in folding of high-flux substrates [26]. However, further information about the specific roles of BamB, BamC and BamE are lacking [21].

In this study, we provide evidence for specialised functions and show that changes to LPS structure lead to increased membrane fluidity and decreased Bam complex activity. We also identify that the cyclic form of enterobacterial common antigen (ECA), a periplasmic carbohydrate, is essential in the absence of the chaperone protein SurA. Lastly, we demonstrate that the Bam accessory lipoproteins, including BamC, are essential in the absence of the peptidoglycan stem-peptide component *meso-*DAP, providing a strong phenotype for strains lacking this poorly understood protein.

## Results

### Loss of Bam-associated proteins leads to distinct phenotypes

We hypothesised that if the Bam-associated proteins have specialised roles in the cell it would be unlikely that they phenocopy each other. Therefore, we tested fitness of Δ*bamB*, Δ*bamC*, Δ*bamE*, Δ*surA*, Δ*skp* and Δ*degP* mutants of *E. coli* K-12 BW25113 compared to the parent strain under various stresses. We arrayed the strains in 384-well format on solid agar plates containing each stress or compound. Endpoint pictures were taken to quantify colony fitness based on size by the image analysis software Iris [27]. Fitness was scored using the chemical genomics analysis software ChemGAPP Small by comparing the mean colony size for the mutant in each condition to the mean colony size of that mutant in the LB agar condition, which was normalised to a fitness score of 1 [28]. Fitness scores below 1 represent decreased fitness, as a function of colony size compared to growth on LB agar, and scores above 1 indicate increased fitness in that condition.

The fitness of each mutant across all conditions identified that the Δ*bamB* and Δ*surA* mutants have the strongest fitness defects **(Fig 1)**. Correlation between each mutant fitness profile pair was assessed by calculating the Pearson correlation coefficient. This demonstrated that Δ*bamB* and Δ*surA* have a similar phenotypic profile with a correlation coefficient of 0.74, the highest of all pairs of mutants **(Fig S1)**. Similar decreased fitness scores were observed for both strains in the presence of bacitracin, carbenicillin, erythromycin, sodium dodecyl-sulfate (SDS), osmotic stress conditions due to varied salt concentrations, and vancomycin **(Fig 1)**. This is unsurprising as SurA is the major chaperone pathway for OMPs to reach the Bam complex resulting in low fitness of Δ*bamB* and Δ*surA* when growing on LB at 37°C. However, they do not phenocopy in all conditions. For example, the Δ*bamB* strain grew better in the presence of the MreB inhibitor A22 with a fitness score of 1.59, but the Δ*surA* mutant was less fit in this condition (score of 0.51). They also had opposing fitness scores in the presence of doxycycline (Δ*bamB* – 1.65, Δ*surA* – 0.94) and hydroxyurea (Δ*bamB* – 0.68, Δ*surA* – 1.14) **(Fig 1)**. The Δ*bamE* mutant mildly correlated to Δ*bamB* with a coefficient of 0.51 **(Fig S1)**. Both mutants were sensitive to vancomycin (Δ*bamB* – 0.06, Δ*bamE* – 0.66) and bacitracin (Δ*bamB* – 0.02, Δ*bamE* – 0.58) two antibiotics that exceed the exclusion limit of outer membrane porins and to which sensitivity indicates damage to the permeability barrier. In contrast, Δ*bamC* has no strong phenotypes and correlated most strongly with the WT parent strain **(Fig 1 and Fig S1**). Similarly to Δ*bamC*, the Δ*skp* mutant also has no strong phenotypes in our screen. The Δ*degP* strain was negatively correlated with the other periplasmic chaperone pathway group mutants Δ*skp* (-0.51) and Δ*surA* (-0.37) and with Δ*bamE* (-0.68) due to opposing fitness scores in a range of stresses **(Fig. 1 and Fig S1)**. Interpretation of this result is complicated by DegP not only functioning as a chaperone, but also as a protease that targets misfolded OMPs in the periplasm [29]. However, due to the role of DegP in degrading stalled OMPs that have been recognised by Skp, they might have been expected to correlate [8]. Together, the phenotypic profiling data indicate that while there are shared phenotypes between each knockout, we observe a broad range of unique, mutant specific phenotypes.

**Figure 1.**
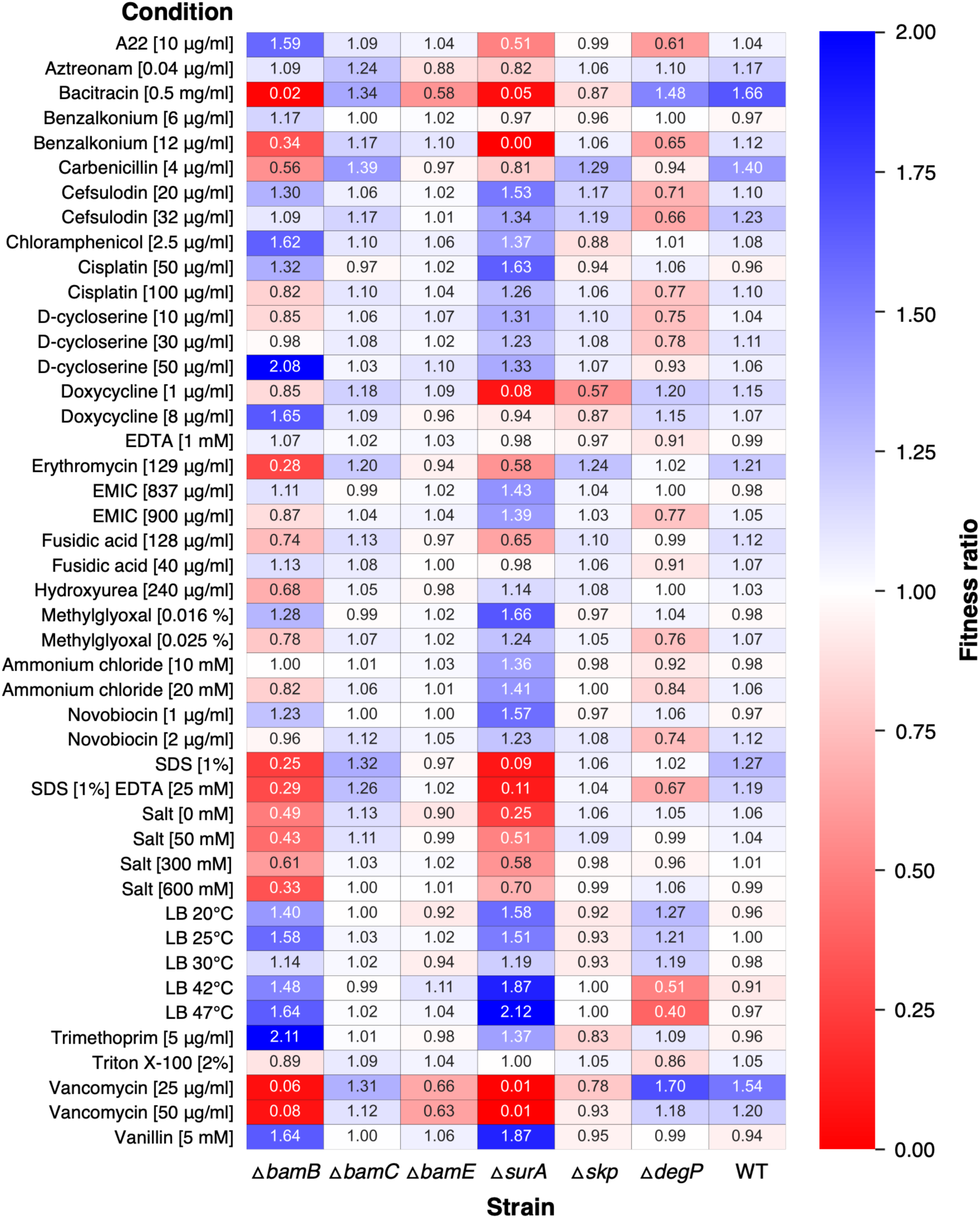
Phenotypic profiling of mutants lacking Bam-associated proteins. Heat map of fitness scores for the *E. coli* BW25113 parent strain and Δ*bamB*, Δ*bamC*, Δ*bamE,* Δ*surA*, Δ*skp*, or Δ*degP* mutants grown in various stress conditions. Colony size measurements for each condition were normalised by comparing to colony size of that strain when grown on LB medium at 37°C. Growth of each strain on LB is set to 1 and each condition normalised to this. Fitness scores above 1 represent better growth than the LB condition as measured by colony size and scores below 1 indicate smaller colony size than the LB condition. Some conditions are abbreviated due to space restrictions: EMIC - 1-Ethyl-3-methylimidazolium chloride, EDTA - Ethylenediaminetetraacetic acid, SDS – Sodium dodecyl sulfate.

### TraDIS identifies unique synthetic-lethal interactions for Bam-associated proteins

Considering that the Bam-associated proteins potentially have specialised function, we sought to find their genetic interaction networks by using Transposon-Directed Insertion site Sequencing (TraDIS). When TraDIS libraries are generated in single gene deletion mutant backgrounds, the insertion frequencies of genes/regions compared to wild-type cells identifies genes that become more important for survival in the mutant backgrounds. These genes are termed synthetic-lethal, and help to identify co-dependencies of processes that the candidate gene might be involved in. The method can also identify mutants that are more fit in the given genetic background or condition [30–33]. Therefore, the parent strain (*E. coli* BW25113) and Δ*bamB,* Δ*bamC,* Δ*bamE,* Δ*surA,* Δ*skp,* or Δ*degP* mutants were transformed with a mini Tn5-kanamycin transposon and recovered on LB agar at 37°C. Transposon mutants were then pooled and transposon insertion sites identified by DNA sequencing as described previously [32]. The BioTraDIS pipeline was used for data analysis to classify genes as either essential or non-essential. Essential genes were then compared to the parent BW25113 TraDIS library to identify genes that became essential in the knockout strains (synthetic-lethal) **(Table S1)** [34]. For quality control, two independent replicates of each transposon library were sequenced over several sequencing runs. Individual sequencing runs had a high correlation coefficient of ≥0.93 between samples and generated >6 million TraDIS reads and >500,000 unique insertion sites for each strain **(Fig S2 and S3)**. Transposon insertion sites were evenly mapped throughout the genome, with the exception of an increased density around the origin of replication as expected due to gene dosage **(Fig S4)** [35].

These experiments are designed to identify novel genetic interactions, but this unbiased approach will also uncover known synthetic-lethal gene pairs that allow benchmarking of our data. We confirmed known genetic interactions between *surA* and *skp* [4], *degP* and *surA* [4], *bamB* and *degP* [26] and *bamE* and *bamB* [25, 36] **(Fig S5)**. The Δ*bamB,* Δ*bamC* and Δ*bamE* TraDIS libraries allowed identification of 93, 92 and 29 candidate synthetic -lethal genes, respectively **(Fig 2A and Table S1)**. This is particularly surprising as there are no known roles or dependencies for *bamC* and we did not observe strong phenotypes for Δ*bamC* in our phenotypic screen **(Fig 1)**. Unfortunately, 54 of the genes identified as synthetic-lethal in the Δ*bamC* background are of unknown function **(Table S1)**. We then compared the candidate synthetic-lethal gene lists to probe if they are involved in similar processes or contribute to the same function within the cell. In the case that all three Bam accessory proteins solely contribute to Bam function, we would expect to find significant overlap between mutants. However, we found many unique synthetic-lethal genes for each (62, 57 and 5 for Δ*bamB,* Δ*bamC* and Δ*bamE,* respectively), supporting the hypothesis that each has specialised roles in the cell **(Fig 2A)**.

**Figure 2.**
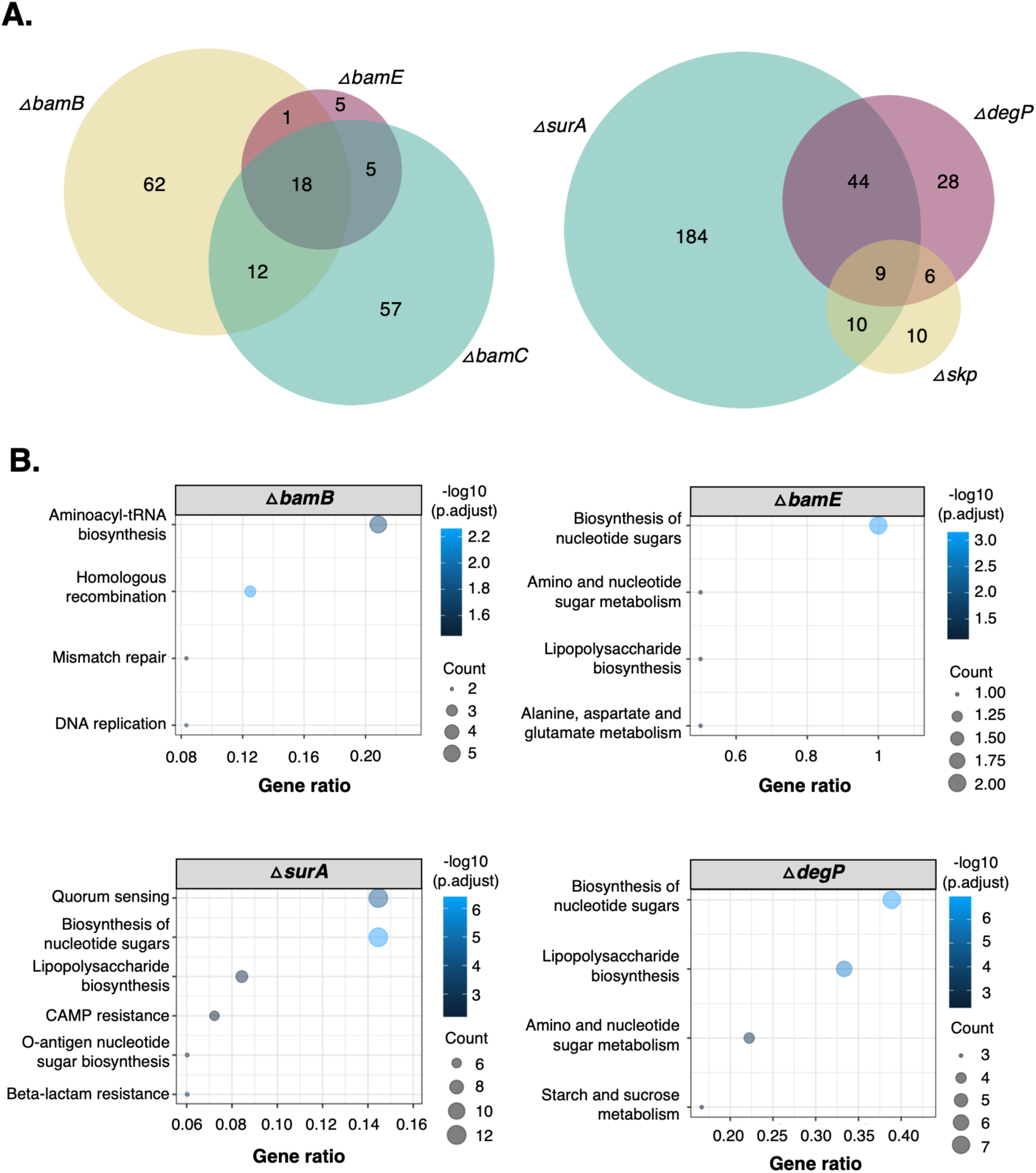
Genetic interaction analysis of Bam-associated genes by TraDIS. TraDIS libraries were constructed in the *E. coli* BW25113 parent strain and Δ*bamB*, Δ*bamC*, Δ*bamE,* Δ*surA*, Δ*skp*, or Δ*degP* mutants to identify genes that become essential in each knockout strain compared to the parent library (synthetic-lethal). **A.** Venn diagrams showing synthetic-lethal genes identified within the Δ*bamB*, Δ*bamC* and Δ*bamE* TraDIS libraries or the Δ*surA*, Δ*skp* and Δ*degP* libraries. **B.** Analysis of KEGG categories enriched in synthetic-lethal gene lists in the Δ*bamB*, Δ*bamE,* Δ*surA* and Δ*degP* TraDIS libraries, represented as bubble plots. Gene ratio represents the proportion of genes within the synthetic-lethal gene set assigned to a given KEGG pathway. No data is shown for the Δ*bamC* and Δ*skp* synthetic-lethal gene lists as there was no significant enrichment of any KEGG categories.

For the OMP chaperone pathway mutants Δ*surA,* Δ*degP* and Δ*skp,* we identified 247, 87 and 35 candidate synthetic-lethal genes, respectively **(Fig 2A)**. The Δ*surA* mutant had many strong phenotypes when probed against stresses **(Fig 1)**, which could explain the large number of identified genes compared to the other OMP chaperone mutants **(Fig 2A)**. Of the 87 candidate synthetic-lethal genes in the Δ*degP* dataset, only 15 were also in the Δ*skp* background, despite DegP and Skp functioning together to degrade stalled OMP substrates [8]. This difference is also potentially due to the dual role of DegP as both a chaperone and a protease [29]. These results suggest that while there is functional redundancy between the SurA and DegP/Skp chaperone pathways, the function of the chaperones under specific chemical and genetic conditions is likely specialised **(Fig 1 and 2A)**. To determine the functions and associated pathways of the candidate synthetic-lethal genes identified in the mutant backgrounds, we completed GO and KEGG analyses. Enrichment analysis of KEGG pathways for the Δ*bamC* and Δ*skp* datasets resulted in no significant enrichment of any one category. However, genes involved in “Lipopolysaccharide biosynthesis” and “Biosynthesis of nucleotide sugars” were enriched in the Δ*surA*, Δ*degP* and Δ*bamE* candidate synthetic-lethal gene lists and “O-antigen nucleotide sugar biosynthesis” was enriched in the Δ*surA* background **(Fig 2B and Table S2)**.

The most enriched KEGG pathway in the Δ*bamB* dataset was “Aminoacyl-tRNA biosynthesis”, however these genes all encode tRNAs and are on average 75 bp in length. Visual inspection of these gene regions identified sparse insertion density within the local genomic area leading to a likely false positive for essentiality. The Δ*bamB* dataset is also significantly enriched for three other KEGG pathways: “Homologous recombination”, “Mismatch repair” and “DNA replication” **(Fig 2B and Table S2)**.

Considering the potential for polar effects in the Δ*bamB* strain on the essential gene coded downstream of *bamB* on the chromosome: *der*, which encodes an essential GTPase involved with ribosomal stability, we sought to validate this surprising result **(Fig S6A)**. Within this set of candidate synthetic-lethal genes, we chose to focus on those with the clearest essentiality from the TraDIS data, *hda* and *seqA.* Insertions within *seqA* and *hda*, which encode negative regulators of DNA replication initiation [37–41], led to decreased survival of Δ*bamB* **(Fig S6)**. We confirmed the findings regarding the negative regulators by using CRISPRi. The *bamB::aph* allele was transferred into the dCas9 encoding strain *E. coli* LC-E18[42] and the kanamycin resistance marker was removed. The pSGRNA plasmid was used to express guide RNA targeting the *seqA* or *hda* genes in either the parent or Δ*bamB* strains carrying pBAD plasmids encoding either *bamB* or *der*. Expression of guide RNA targeting the *seqA* and *hda* genes led to significantly decreased survival of Δ*bamB* when compared to the parent strain, therefore confirming the observations made from TraDIS. However, expression of *bamB* from the pBAD plasmid did not complement this effect, but expression of *der* did, therefore confirming the genetic link to DNA replication was caused by effects on expression of the *der* gene in the Δ*bamB* strain **(Fig S7)**. We also confirmed that the Δ*bamB* strain has disrupted DNA replication initiation control by completing replication run-out assays[43]. Our TraDIS data and double mutant analysis also confirmed that disruption of the positive regulator *diaA* in the Δ*bamB* strain increases the strain fitness **(Fig S8-12)**. While not within the scope of this study, the effects caused by *der* gene expression perturbation will likely be of interest to researchers of this essential GTPase. Despite this complication, the KEGG pathway enrichment analysis identifies the importance of LPS and ECA biosynthesis in multiple mutants reducing the likelihood of complications from local effects of gene knockout construction. Therefore, this directed our further investigations.

### Genes required for heptose incorporation into LPS are synthetically lethal with *bamB, surA* and *degP*

We identified that fewer mutants were recovered with transposon insertions in genes involved in LPS assembly in the Δ*bamB*, Δ*surA* and Δ*degP* backgrounds than in the parent. In addition, genes involved in “Biosynthesis of nucleotide sugars” were significantly enriched, with genes identified being involved in the synthesis of heptose, a core component of LPS **(Fig 2B and Table S2)**. Despite this category being enriched in the Δ*bamE* dataset, we found this was due to one gene in particular, *glmS*, and that the result was specific to the Δ*bamB*, Δ*surA* and Δ*degP* TraDIS libraries **(Fig 3A and Fig S13)**. Synthesis of heptose consists of five main steps before it is incorporated into the LPS inner core **(Fig 3B)**. In the Δ*surA* and Δ*degP* TraDIS libraries, all four genes involved in synthesis of heptose were essential: *gmhA*, *gmhB*, *hldE* and *waaD*. In the Δ*bamB* background all were essential except for *gmhB,* which is potentially because a Δ*gmhB* mutant is not completely devoid of heptose [44]. These data suggest that heptose production was functionally more important in Δ*degP* and Δ*surA* than in the Δ*bamB* mutant. Incorporation of heptose into the LPS structure was also functionally important. In Δ*degP* and Δ*surA* mutants, the genes *waaC* and *waaF* were synthetic-lethal, which are responsible for transfer of the first and second heptose onto the LPS inner core, respectively **(Fig 3A and 3B)** [45, 46]. In contrast, in the Δ*bamB* mutant, *waaC* was synthetic-lethal whereas *waaF* was not. This implies that while Δ*degP* and Δ*surA* mutants are unable to tolerate having LPS with only one heptose residue, the Δ*bamB* mutant is.

**Figure 3.**
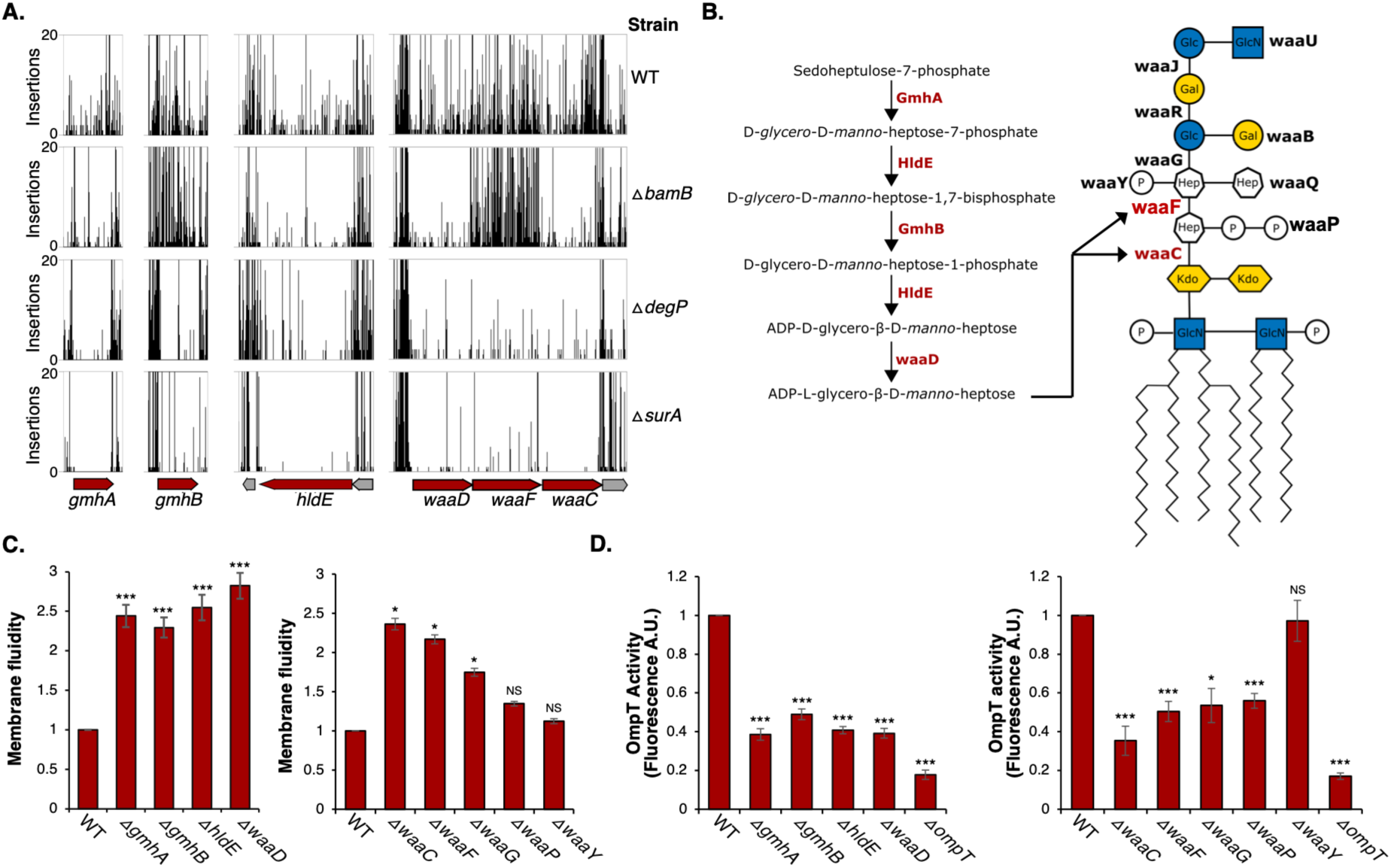
Incorporation of heptose into LPS is essential in Δ*bamB,* Δ*surA* and Δ*degP*. **A.** Transposon insertions in the genes *gmhA*, *gmhB*, *hldE, waaD, waaC,* and *waaF* in the parent, Δ*bamB,* Δ*surA* and Δ*degP* TraDIS libraries. Transposon cut-off is set to 20. Essential genes are represented as red arrows. **B**. Schematic of the heptose biosynthesis pathway and LPS structure in *E. coli* K-12 BW25113 with gene labels for LPS biosynthesis enzymes indicated next to the linkage they form or component they ligate. Enzymes identified as synthetic-lethal are labelled in red text. **C.** Membrane fluidity was measured in Δ*gmhA,* Δ*gmhB,* Δ*hldE,* Δ*waaD,* Δ*waaC*, Δ*waaF*, Δ*waaG*, Δ*waaP* and Δ*waaY* mutants and compared to the parent strain by using pyrenedeconoic acid fluorescence. **D.** To determine the extent of Bam complex activity, successful folding and insertion of OmpT into the outer membrane was measured in the Δ*gmhA,* Δ*gmhB,* Δ*hldE,* Δ*waaD,* Δ*waaC*, Δ*waaF*, Δ*waaG*, Δ*waaP*, Δ*waaY* and Δ*ompT* mutants and compared to the parent strain. For panels C and D, experiments were performed in biological and technical triplicate with standard deviation represented by error bars. Two sample t-tests were used to assess statistical significance of differences from the WT strain with *** indicating p-values of <0.001, * indicating p-values of <0.05, and NS as not significant (p-value ≥0.05).

### LPS truncation increases membrane fluidity and decreases Bam complex activity

Increased outer membrane fluidity decreases Bam activity and modifications to LPS structure can lead to changes in membrane fluidity [47]. Thus, we hypothesised that changes in LPS core structure would affect membrane fluidity to differing degrees, which could affect Bam complex efficiency. To test this hypothesis, we assessed membrane fluidity and Bam complex activity in strains lacking the genes encoding the heptose biosynthesis pathway (Δ*gmhA*, Δ*gmhB,* Δ*hldE*, and Δ*waaD*) and LPS core synthesis (Δ*waaC*, Δ*waaF*, Δ*waaP*, Δ*waaG* and Δ*waaY*) **(Fig 3)**. There were no differences in transposon insertion index for the gene *waaY,* which encodes the enzyme responsible for phosphorylation of the second heptose in the LPS inner core. Therefore, Δ*waaY* should act as a control in this experiment [48]. In addition, Δ*waaP* and Δ*waaG*, which are responsible for phosphate addition to the first heptose, and incorporation of the first glucose, respectively [49], were included as a small decrease in the insertion indexes for these genes were observed in the Δ*surA* and Δ*degP* TraDIS datasets **(Fig S14).**

Fluidity of the membrane for each mutant was measured using the lipophilic pyrene probe, pyrene decanoic acid. The pyrene monomer can undergo excimer formation and demonstrates a shift in fluorescence, a process which is dependent on the ease of mobility within the membrane [50, 51]. Formation of the excimer was measured and compared to that of the parent strain **(Fig 3C)**. The largest increase in membrane fluidity occurred in knockouts of genes required for heptose biosynthesis: Δ*gmhA*, Δ*gmhB,* Δ*hldE*, and Δ*waaD*. Of this group, the Δ*gmhB* mutant had the smallest increase in membrane fluidity and this gene was also not essential in the Δ*bamB* background **(Fig 3A and 3C)**. Of the genes required for LPS core biosynthesis, membrane fluidity was higher in Δ*waaC*, Δ*waaF* and Δ*waaG* than in the parent strain with the biggest change being in Δ*waaC* and the smallest in Δ*waaG*. However, there was no significant increase in fluidity in the Δ*waaP* mutant. Membrane fluidity of the Δ*waaF*, Δ*waaP* and Δ*waaG* mutants was intermediate between that of the heptoseless Δ*waaC* mutant and the parent, with the severity of the effect correlating to the severity of LPS core truncation. No significant change in membrane fluidity was measured in the Δ*waaY* mutant control **(Fig 3C)**.

Next, we investigated whether impairment of heptose synthesis affected activity of the Bam complex, which was monitored by an *in vivo* OmpT fluorescence assay. The outer membrane protease OmpT requires the Bam complex for folding and insertion into the membrane. Folded OmpT is able to cleave a fluorogenic peptide, which is monitored by increased fluorescence over time [10]. Except for Δ*waaY*, all mutants demonstrated a minimum 40% decrease in OmpT activity compared to the parent strain. The heptoseless Δ*waaC* mutant exhibited the greatest decrease in OmpT activity. The effect of Δ*waaF*, Δ*waaP* and Δ*waaG* mutations on OmpT activity levels was comparable to each other and intermediate between that of Δ*waaC* and the parent strain **(Fig 3D)**. In summary, mutations that lead to heptoseless LPS had the biggest increase in membrane fluidity and the lowest levels of OmpT activity with a graded response based on the severity of LPS core truncations. This suggests there is a correlation between LPS core structure, membrane fluidity and Bam complex activity that leads to a variety of Bam complex activity levels depending on the form of LPS in the outer membrane.

### Generation of cyclic ECA is essential in the absence of the chaperone SurA

Genes involved in ‘Biosynthesis of nucleotide sugars’ were significantly enriched as synthetic-lethal in the Δ*surA* mutant. The genes identified are involved in biosynthesis of ECA (**Fig 2B)**, which is a highly conserved carbohydrate-derived molecule present on the external leaflet of the outer membrane and in the periplasm of Enterobacteriaceae [52, 53]. The ECA biosynthesis pathway genes were all synthetic-lethal, with the exception of *wxyE,* which is required for survival of both the parent and the Δ*surA* mutant **(Fig 4A and 4B)**. Also, fewer mutants were recovered with transposon insertions in the genes *rffH* and *rffG* than in the parent TraDIS library (**Fig 4A)**. The gene products RffH and RffG catalyse the same enzymatic reaction and are homologous in sequence to the genes RfbA and RfbB, respectively. However, they form part of different operons and function in separate pathways despite *rffG* being able to complement an RfbB defective strain [54, 55]. This likely explains why *rffH* and *rffG* are not entirely essential in the Δ*surA* background.

**Figure 4.**
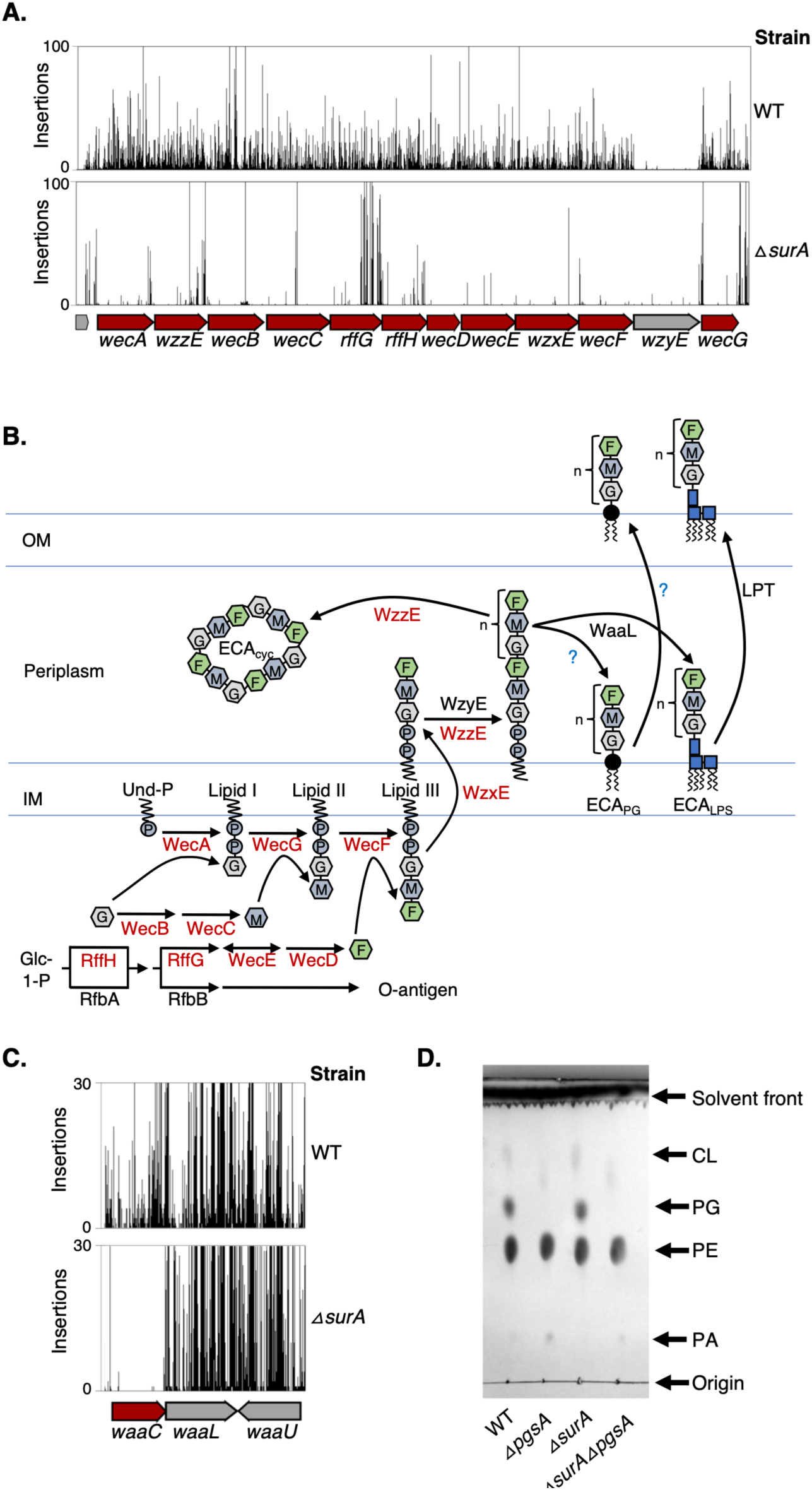
Synthesis of ECA is essential in the absence of the chaperone SurA. **A.** Transposon insertions in genes of the ECA biosynthesis pathway in the parent or Δ*surA* mutant TraDIS libraries. Transposon cut-off is set to 100. Essential genes are represented as red arrows. **B**. Schematic representation of the ECA biosynthesis pathway with the names of proteins labelled next to the reaction for which they are responsible. Synthetic-lethal proteins are labelled in red text. **C.** Transposon insertions in the *waaL* gene in either the parent or Δ*surA* mutant TraDIS libraries. Transposon cut-off is set to 100. **D.** Analysis of phospholipid content in the parent or Δ*surA* mutant with further mutations to disrupt synthesis of the major anionic phospholipids. The Δ*pgsA and* Δ*surA*Δ*pgsA* strains are also Δ*lpp*Δ*rcsF*. Phospholipid samples were separated and visualised by TLC using a solvent mixture of methanol/ chloroform/ water with a ratio of 2:2:1.8 before being visualised by phosphomolybdic acid and charring.

ECA exists in three forms that all share the same biosynthetic pathway: ECA that is covalently linked to LPS (ECA_LPS_), covalently linked to diacylglycerol-phosphate (ECA_PG_), or a cyclic form (ECA_CYC_) that is localised to the periplasm as opposed to being surface exposed [52, 56]. We sought to determine which form of ECA is synthetic-lethal in Δ*surA*. Synthesis of ECA_LPS_ is facilitated by the O-antigen ligase WaaL, which is responsible for attaching the ECA molecule onto the LPS core [57, 58]. However, in the Δ*surA* TraDIS library, the gene *waaL* is non-essential **(Fig 4C)**. This suggests that ECA_LPS_ is not essential in the Δ*surA* mutant. The formation of ECA_PG_ is completed by attachment of linear ECA chains to diacylglycerol by a phosphodiester bond through an unknown mechanism [57, 59]. To determine the essentiality of ECA_PG_ we targeted synthesis of the donor molecule for ECA_PG_, the phospholipid phosphatidylglycerol [60]. The gene *pgsA* encodes phosphatidylglycerophosphate synthase, which catalyses the first committed step in biosynthesis of phosphatidylglycerol. However, loss of *pgsA* is lethal due to mislocalisation of Braun’s lipoprotein, Lpp, to the inner membrane and activation of the Rcs stress system. These issues can be resolved by making an Δ*lpp*Δ*rcsF* mutant before constructing the Δ*pgsA* mutation [61–63]. The Δ*pgsA*Δ*surA*Δ*lpp*Δ*rcsF* quadruple mutant was viably constructed and the absence of phosphatidylglycerol confirmed by phospholipid extraction and thin layer chromatography **(Fig 4D)**. This confirmed that the phospholipid donor for synthesis of ECA_PG_ is not essential in the Δ*surA* mutant. Lastly, the gene *wzzE* is not required for production of ECA_LPS_ or ECA_PG_, but is required for production of ECA_CYC_ [57, 64, 65]. In the Δ*surA* TraDIS library, the gene *wzzE* contained an essential region indicating that ECA_CYC_ is likely to be the form of ECA that is essential in a Δ*surA* mutant **(Fig 4A)** [64].

### *meso*-DAP is essential in the absence of BamB, BamC or BamE

The TraDIS data were searched for synthetic-lethal genes involved in cell envelope biogenesis pathways other than LPS and ECA biosynthesis. The gene *dapF* was identified as essential in the Δ*bamB* and Δ*bamC* strains by visual inspection indicating decreased insertions in *dapF* in the Δ*bamE* strain **(Fig 5A)**. This was particularly surprising considering the lack of strong phenotypes in the Δ*bamC* background **(Fig 1)**. DapF converts LL-diaminopimelate (LL-DAP) to *meso*- diaminopimelate (*meso*-DAP), which is then either decarboxylated to produce L-lysine by the enzyme LysA or is used in biosynthesis of peptidoglycan [66, 67]. The gene *lysA* was non-essential in all strains tested, suggesting the essentiality of *dapF* in these mutants is due to the requirement for *meso*-DAP in peptidoglycan **(Fig 5A and 5B)**. To validate the genetic interaction between Δ*dapF* and components of the Bam complex, the *dapF* gene was knocked out in the BW25113 parent strain, Δ*bamB,* Δ*bamC* or Δ*bamE* mutants. The double knockouts were selected in the presence of externally supplied *meso*-DAP, which alleviated the loss of *dapF*. Cells were then grown in liquid media in the presence of *meso*-DAP before being serially diluted in un-supplemented media and assayed for survival in the presence or absence of *meso*-DAP by efficiency of plating assay. All strains grew equally as well as the parent on LB agar supplemented with 1 mM *meso-*DAP, except for Δ*bamB*Δ*dapF* which showed a minor decrease in CFU and colony size **(Fig 5C)**. However, in the absence of *meso*-DAP the single Δ*dapF* mutant demonstrated a decrease in survival and the double mutants were not viable **(Fig 5C)**. Lastly, considering that loss of DapF will lead to a weaker peptidoglycan layer due to decreased crosslinks, we sought to negate the synthetic-lethal phenotype by growth in the presence of sucrose as an osmoprotectant [68, 69]. Unexpectedly, while the presence of 10% sucrose enabled survival of the single *dapF* mutant to levels comparable to the parent, this was insufficient to restore growth of the double mutant **(Fig 5C)**. This data demonstrates that peptidoglycan structure is of increased importance in the absence of full Bam complex activity and that this may not simply be due to structural support for the envelope.

**Figure 5.**
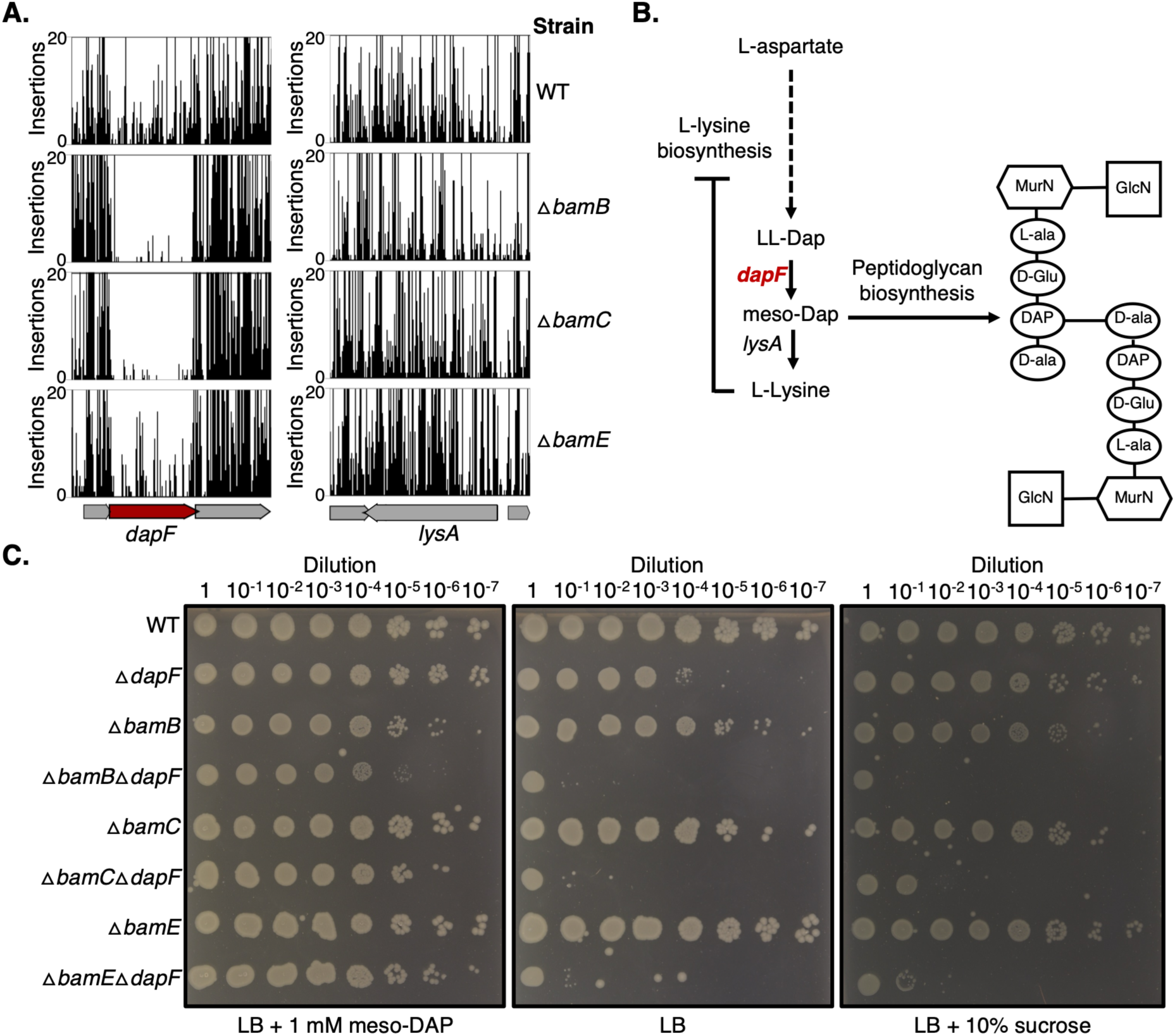
*meso*-DAP containing peptidoglycan is essential in the absence of Bam accessory lipoproteins. **A.** Transposon insertions in the genes *dapF* and *lysA* in the parent or Δ*bamB,* Δ*bamC,* Δ*bamE* mutant TraDIS libraries. Transposon cut-off is set to 20. Essential genes are represented as red arrows. **B**. Schematic representation of the L-lysine biosynthesis pathway with the names of proteins labelled next to the reaction for which they are responsible. Synthetic-lethal proteins are labelled in red text. A schematic structure of *E. coli* peptidoglycan is shown with the site of *meso*-DAP incorporation shown. **C.** Efficiency of plating assay showing survival of the Δ*dapF,* Δ*bamB,* Δ*bamC,* Δ*bamE,* Δ*dapF*Δ*bamB,* Δ*dapF*Δ*bamC, or* Δ*dapF*Δ*bamE* mutants grown on LB with or without 1 mM *meso*-DAP or 10 % sucrose. Cells were grown overnight in LB supplemented with 1 mM *meso*DAP before being normalised to OD_600_ = 1.00 and serially diluted 1:10 before 2 μl was spotted on agar plates and incubated at 37°C overnight.

### OMP trafficking protein conservation varies throughout Gram-negative bacteria

The data presented here demonstrate that in *E. coli,* the Bam-associated proteins likely have specialised functions that are potentially coordinated with LPS structure, the presence and form of ECA and peptidoglycan biogenesis. Considering these results, we revisited the evolutionary conservation of Bam complex subunits and the periplasmic chaperone pathway proteins [70]. Reference genomes for a diverse range of Gram-negative bacterial representative strains were used to search for sequences coding the Bam complex subunits (as observed in *E. coli*) or the chaperone pathway proteins SurA, Skp and DegP. A combination of Prokka annotation, hmmsearch, SignalP 6.0 [71] and manual curation were used to generate a neighbour joining tree with a heat map of gene counts in each organism **(Fig 7)**. While we found broad conservation of all query proteins within the gamma- and beta-proteobacteria, there are some notable exceptions. The BamB and BamC lipoproteins are not conserved within two species of the aphid endosymbiont *Buchnera aphidicola* and the data confirm previous unpublished observations that a strain of *B. aphidicola*, (subspecies *Baizongia pistaciae*) appears to have entirely lost genes encoding the Bam complex [72]. This is possibly due to the strain containing only a few OMPs, therefore no longer requiring a dedicated OMP assembly machinery.

**Figure 7.**
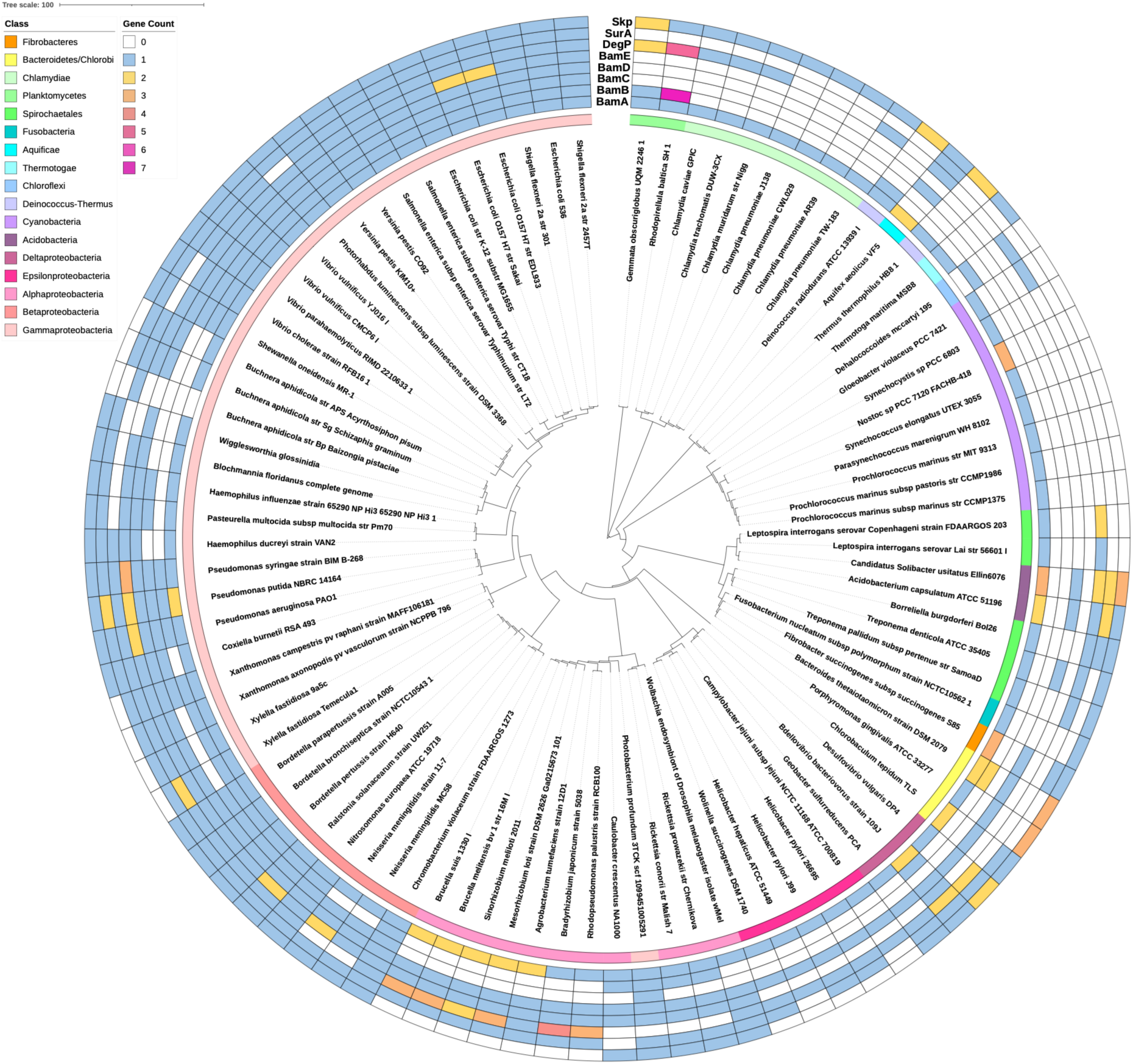
Conservation analysis of Bam-associated proteins in Gram-negative bacteria. Conservation of the BamB, BamC, BamE, SurA, DegP and Skp proteins among key Gram-negative bacterial species represented by a presence/absence map plotted onto a phylogenetic tree generated with iTOL [73]. Gene count in each species is represented by coloured boxes.

Outside of the gamma- and beta-proteobacteria, requirement for the Bam-associated proteins appears to be diverse. The lipoproteins BamB and BamC are largely absent in the alpha-proteobacteria, and the epsilon-proteobacteria only encode the core components BamA and BamD, demonstrating reductive evolution. This minimalist form of the Bam complex is sufficient for OMP insertion, which is likely due to reduced β-barrel complexity as observed in *Helicobacter pylori* [74] **(Fig 7)**. While there is diversity in the presence or absence of these proteins in the organisms analysed, multiple copies of some components were also identified. We found that *Pseudomonas putida* encodes two copies of BamA, BamE and SurA. Outside of the gamma-proteobacteria there are numerous examples of species encoding multiple copies of BamA, particularly within the alpha-proteobacteria, such as the plant pathogen *Agrobacterium tumefaciens* **(Fig 7)**. These copies may play specialised roles in folding and inserting specific cargo under specific conditions, they may alleviate an increased requirement for OMP insertion in these organisms or could be required for resistance to Bam-targeting antimicrobials.

## Discussion

The Bam complex has been identified as a significant unrealised target for new antimicrobials, with several new inhibitors of this essential outer membrane biogenesis machine being identified recently [75–78]. However, despite this focus as an important new avenue for drug discovery, and recent advances in understanding the mechanism by which the Bam complex folds and inserts OMPs into the outer membrane [79, 80], we still do not fully understand the roles of the Bam-associated proteins in *E. coli*. We also do not understand why the complex appears to be modular, with subunits being differentially conserved throughout Gram-negative bacteria as shown through our analysis and those done previously **(Fig 7)** [70]. Constituent parts of the outer membrane such as phospholipid species, type of LPS, O-antigen, ECA, lipoprotein content, and variety/flux of the unfolded OMP proteins can vary across bacterial species [81, 82]. Such variation would likely result in significant differences in the biophysical constraints of the membrane environment in which the Bam complex must operate. In addition, there is likely to be differing requirements for the chaperone pathway proteins depending on the pool of client OMPs. Considering that Skp and SurA have been shown to act on different unfolded states of OmpC, with Skp likely being more important during stress conditions, the environmental conditions experienced by each species could also dictate the requirement for each OMP chaperone [6]. Together, this strongly suggests that each accessory lipoprotein is likely to have specialised functions and that the chaperone proteins are unlikely to be truly functionally redundant. This is supported by our phenotypic profiling of knockout strains. Should each of the accessory lipoproteins merely contribute to overall activity of the Bam complex, and if there is true redundancy in the chaperone pathway, then we would expect that the knockout strains should phenocopy each other. However, we found this not to be the case **(Fig 1)**. Our TraDIS analysis also supports this conclusion as we saw many unique synthetic-lethal genes for each mutant. However, validation of this large number of mutants was impractical for this study, but we expect this list of genes will be a useful resource for the research community to develop hypotheses and begin further investigations **(Fig 2)**.

### Role of the outer membrane lipid environment for OMP insertion

The Bam complex must function within the constraints of the outer membrane lipid environment, therefore OMP biogenesis is likely to be affected by changes to outer membrane lipids. Indeed, we found synthetic-lethality between genes involved in LPS inner core biogenesis and either *bamB*, *surA* or *degP*. Complete truncation of the LPS inner core by disruption of the last enzyme in the heptose biosynthesis pathway has previously been demonstrated to increase membrane fluidity and lead to decreased Bam complex activity [47]. We confirmed this observation and further demonstrate that the changes to membrane fluidity are of graded severity relating to the severity of LPS inner core truncation. This in turn leads to a graded impact on Bam complex activity **(Fig 3)**. The Δ*bamB,* Δ*surA* and Δ*degP* knockout strains have the most severe phenotypes within the set of strains tested here (**Fig 1)**. Also, loss of BamB, SurA or DegP each leads to OMP assembly defects, decreased numbers of folded OMPs in the outer membrane, and accumulation of unfolded OMPs in the periplasm [9, 26, 83–85]. We hypothesise that the negative effect of increased membrane fluidity on the Bam complex in combination with decreased levels of OMP assembly in the absence of BamB, SurA or DegP leads to OMP assembly levels that are too low to sustain cell viability. This is of particular significance when considering the variation in LPS structure seen between different strains, and the LPS modifications available, which could affect the level of Bam activity and influence the conservation of each Bam-associated protein [81, 82].

### A role for ECA_cyc_ in amelioration of periplasmic protein folding stress?

The carbohydrate antigen ECA consists of three repeating sugars, is widely conserved amongst the *Enterobacterales*, and is produced in three forms that are either surface exposed (ECA_LPS_ and ECA_PG_) or periplasmically localised (ECA_CYC_). The invariant nature of the antigen suggests it has an important undiscovered function, however the role of ECA in outer membrane biology remains unknown [56]. We identified that ECA biogenesis becomes essential in the absence of the chaperone SurA **(Fig 4A)**. Production of ECA requires the lipid carrier undecaprenyl phosphate (Und-P), which is limited in the cell and utilised in numerous metabolic reactions including the production of peptidoglycan, capsule, O-antigen and membrane-derived oligosaccharides. Disruption of ECA biogenesis can cause stress on these other pathways due to Und-P sequestration in ECA dead-end intermediates [56, 64, 86]. However, the first gene in the ECA pathway is synthetic-lethal in the absence of SurA **(Fig 4A)**, therefore the effects are unlikely due to stress on the Und-P pool. We showed that the surface exposed forms, ECA_LPS_ and ECA_PG_, are not essential in the absence of SurA, suggesting that the periplasmically localised form, ECA_cyc_, is essential **(Fig 4)**. It is possible that ECA_cyc_ might help stabilise the outer membrane of a Δ*surA* mutant, as it has been shown to suppress the envelope permeability defect of a Δ*yhdP* mutant. However, the molecular details of this suppression are yet to be discovered [64]. Alternatively, ECA_cyc_ could be performing a chaperone function directly, but this would require further investigations that are beyond the scope of our study.

### Coordination of OMP biogenesis with peptidoglycan biosynthesis

To ensure successful cell growth and division, activity of the Bam complex must be coordinated with biogenesis of the peptidoglycan layer. Our TraDIS analysis identified that in the Δ*bamB,*Δ*bamC* and Δ*bamE* mutants, the gene *dapF* was essential **(Fig 5A)**. The DapF enzyme is responsible for synthesis of the peptidoglycan stem-peptide component *meso*-DAP. The single and double mutants could be rescued by exogenous *meso*-DAP, however only the single *dapF* mutant could be rescued by the osmo-protectant sucrose, which is sufficient to allow survival during disruption of peptidoglycan biogenesis **(Fig 5B)** [68]. Loss of *meso*-DAP in the single *dapF* mutant leads to accumulation of LL-DAP, which is incorporated into peptidoglycan and leads to decreased crosslinking [66, 69]. In the Δ*dapF* mutant the peptidoglycan layer would be able to withstand less mechanical load due to decreased crosslinking. This decreased mechanical strength, in combination with the reduced capacity for load bearing by the outer membrane in the Bam complex mutants, could mean that the cells are unlikely to withstand osmotic stress, therefore this could be an effect of the combination of stresses on the envelope [87]. However, while this may be true for Δ*bamB* and Δ*bamE* mutants, there are no strong phenotypic effects for Δ*bamC* (**Fig 1)** [88]. Also, the loss of BamC has only a slight effect on the membrane permeability barrier, no detectable changes to the outer membrane proteome, and no effect on OMP folding in an *in vitro* assay [9, 89]. Consequently, the synthetic lethality between *dapF* and *bamC* is unlikely to be due to osmotic stress tolerance, as could be the case for the other mutants.

It has been demonstrated that the Bam complex preferentially inserts OMPs at the cell division site and that this is coordinated by interaction with the peptidoglycan layer [90]. While all of the Bam complex proteins were shown to interact with peptidoglycan *in vitro*, only BamA and BamC were found to interact with peptidoglycan in whole cells when analysed by crosslinking and pull-down assays [90]. Considering the interaction of mature peptidoglycan with BamC and the altered peptidoglycan structure in the Δ*dapF* strain [66, 69], we hypothesise that BamC might facilitate coordination between the Bam complex and peptidoglycan biosynthesis, however this requires further investigation.

Together, our data demonstrate that the Bam-associated proteins have specialised roles in the cell and highlights potential future targets to understand these roles. We provide further evidence that OMP transport, folding and insertion is affected by, and potentially coordinated with, peptidoglycan and LPS structure.

## Acknowledgments

This work was supported by a UKRI Future Leaders Fellowship [MR/V027204/1], a BBSRC responsive mode grant [BB/Y001265/1] and a Springboard Award [SBF005\1112] to Manuel Banzhaf. The work was also supported by a Royal Society Research Grant to Jack Bryant [RGS\R2\242458] and was funded by the National Science Centre Poland [UMO-2014/13/B/NZ2/01139] (awarded to Monika Glinkowska), the KAUST baseline fund [BAS/1/1108-01-01] awarded to Danesh Moradigaravand who is a member of the KAUST Smart-Health initiative, and the EU ITN Train2Target [721484] that funded training of Kara Staunton.

## Supplementary figures

**Figure S1.**
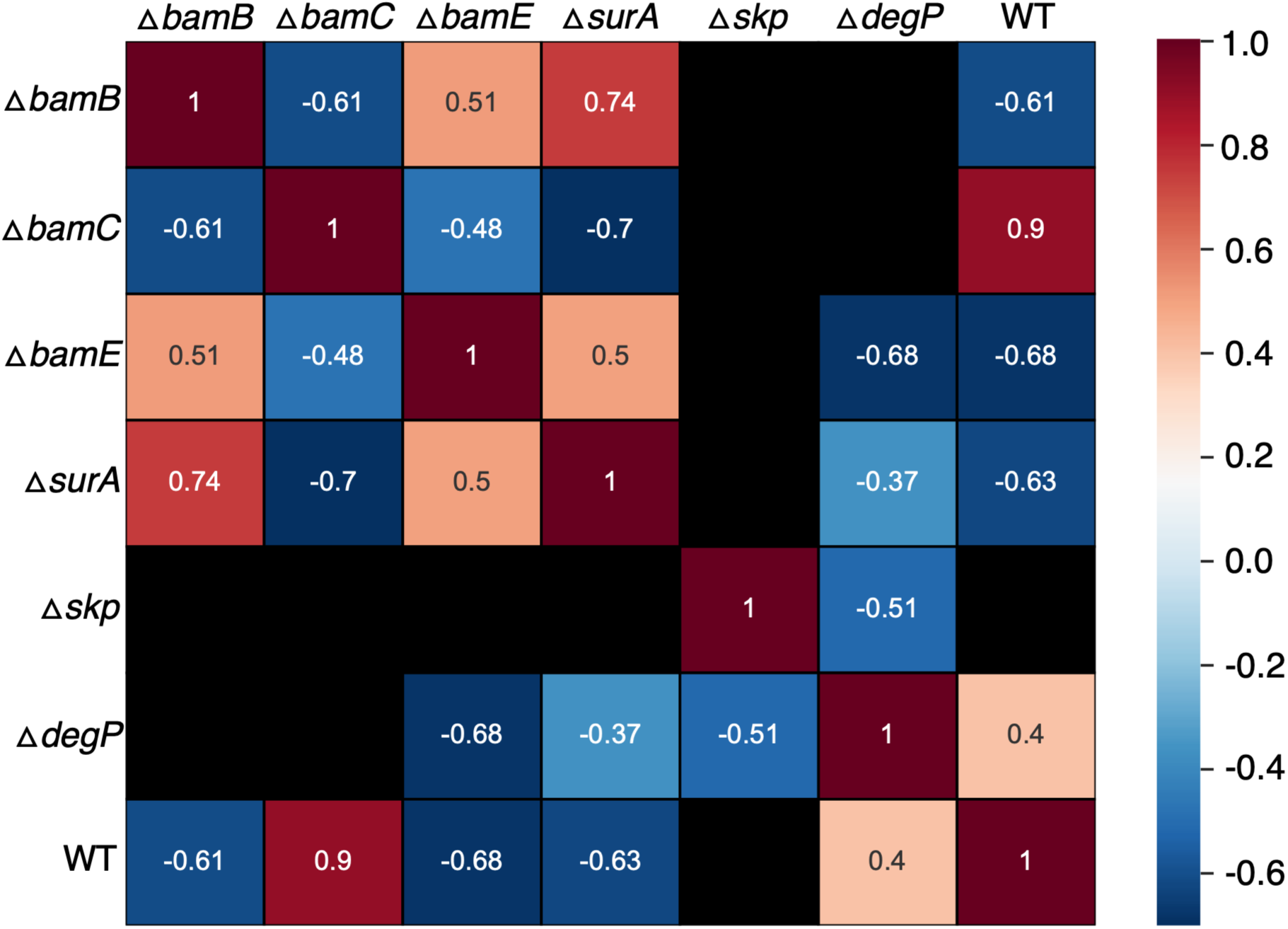
Correlation of phenotypic profiles. Heatmap of Pearson correlation coefficients for each pair of strain phenotypic profiles. Correlation of fitness ratios for each strain was assessed by calculating the Pearson correlation coefficient for averaged fitness scores across replicates using Python before being plotted as a heatmap. Black squares indicate a p-value ≥0.05 meaning the correlation coefficient achieved was not statistically significant.

**Figure S2.**
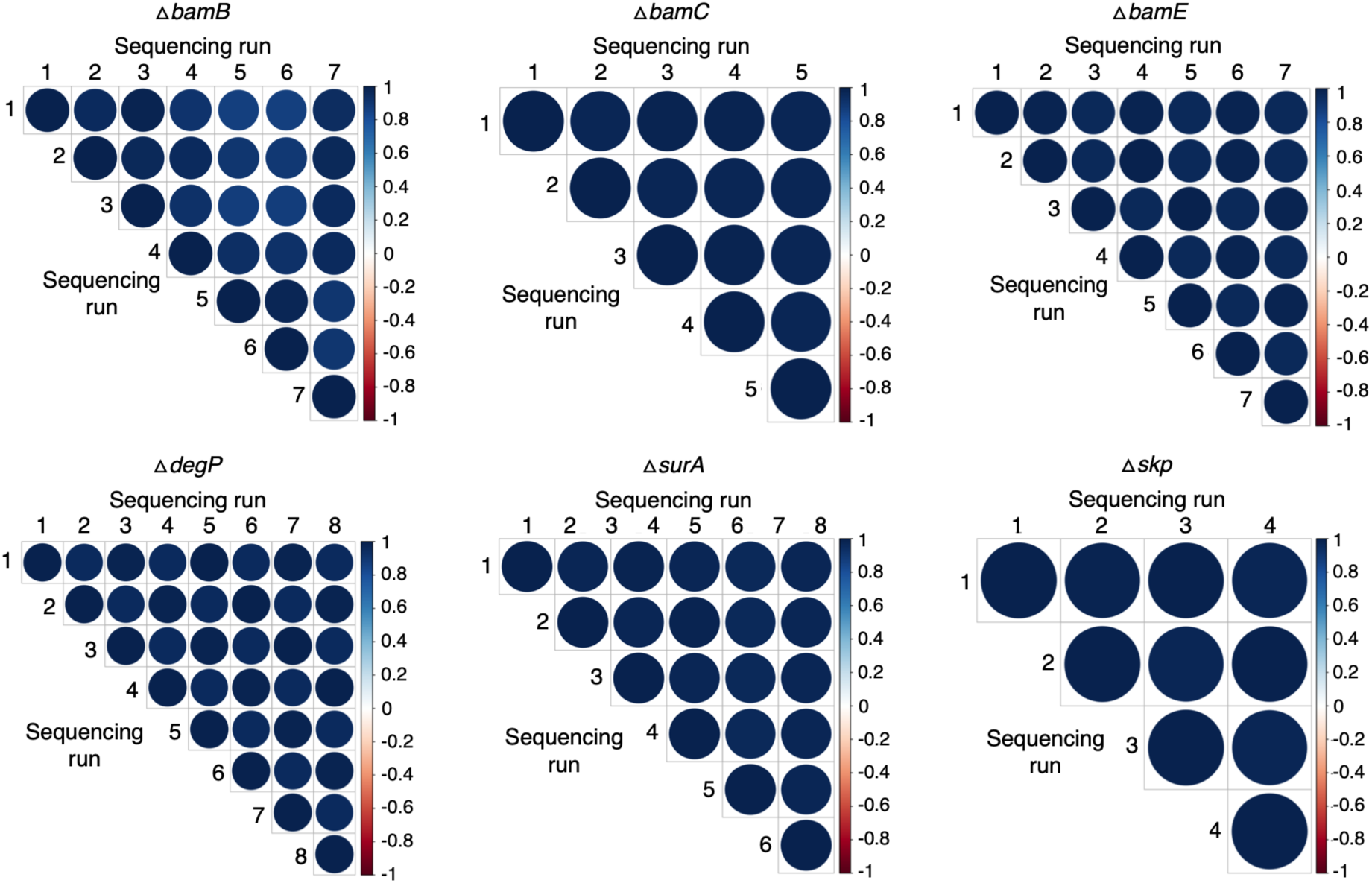
Reproducibility across TraDIS library sequencing runs. Correlogram of insertion indexes between individual sequencing runs for each TraDIS library. The colour and size of the bubbles correspond to the strength of the Pearson correlation coefficient between the indices for the same genes across the runs. The blue colour represents positive correlation.

**Figure S3.**
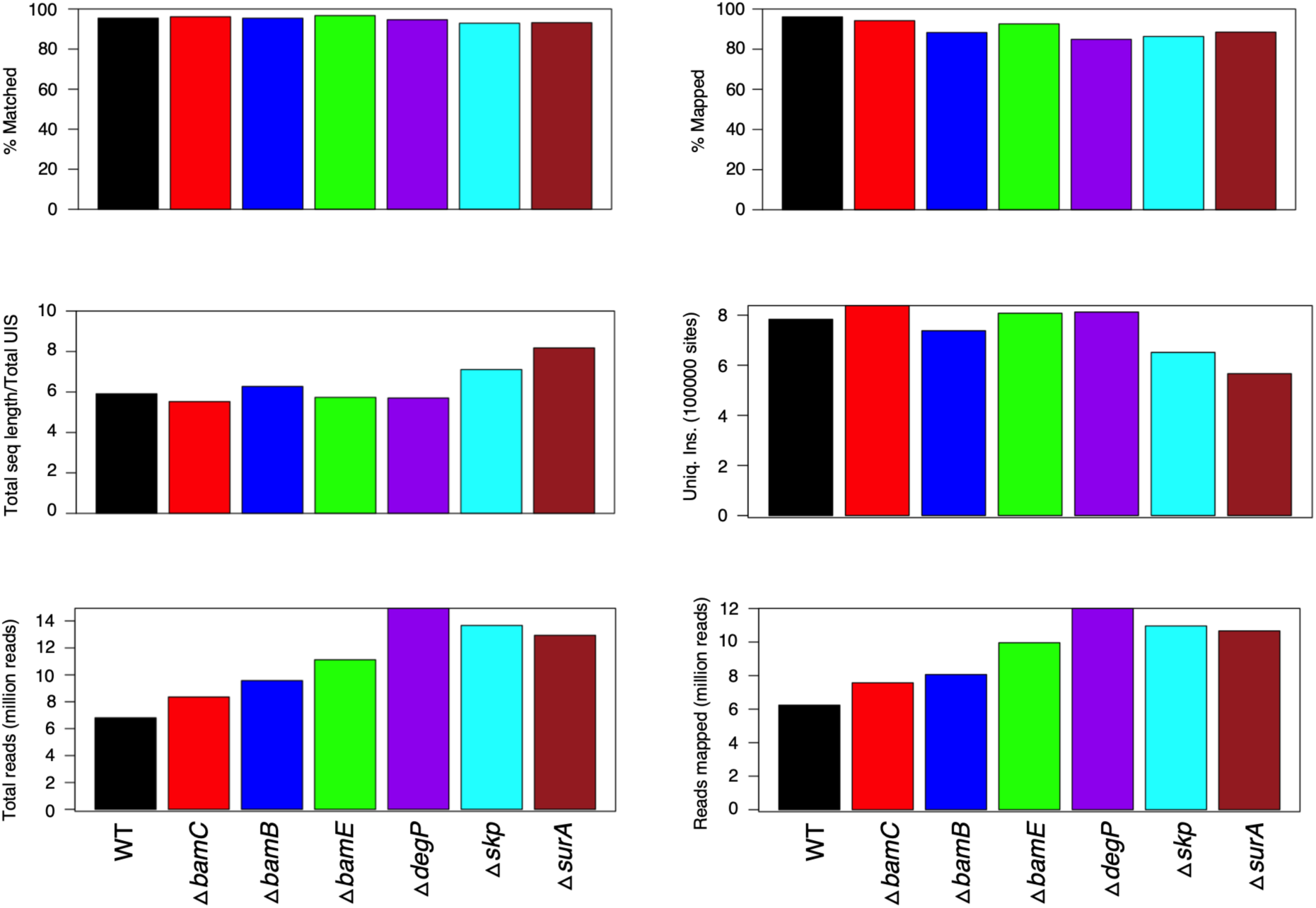
Transposon library construction metrics. TraDIS library construction metrics showing the percentage of sequencing reads that matched the transposon tag and that mapped to the chromosome. Total sequencing length as a function of total unique insertion sites, the total number of unique insertion sites for each sample, total number of sequencing reads, and the number of reads that mapped are also shown.

**Figure S4.**
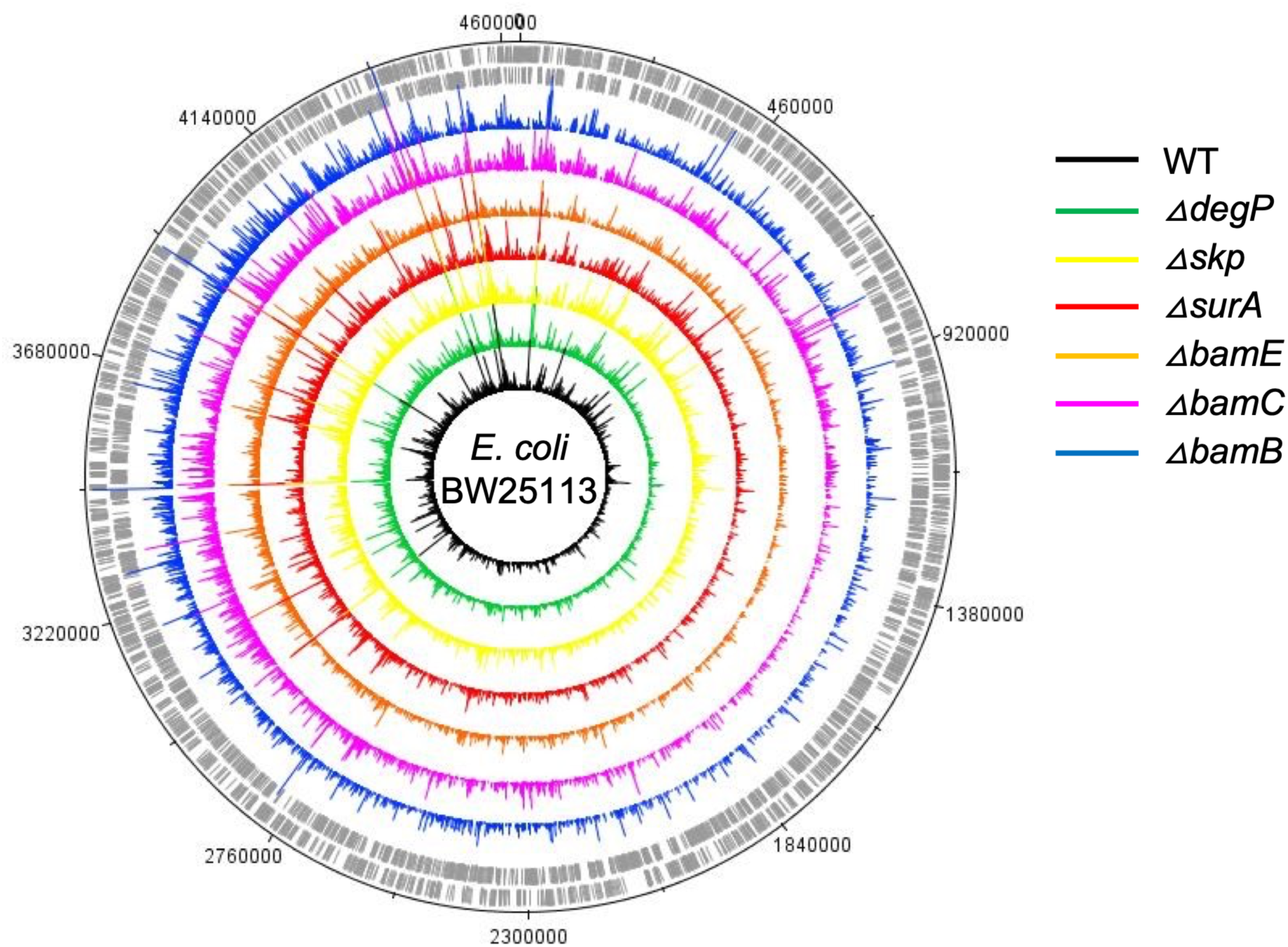
Circular plots of transposon insertion density in TraDIS libraries. Circular plot of transposon insertion sites in the parent BW25113 (WT), Δ*degP,* Δ*skp,* Δ*surA,* Δ*bamE*, Δ*bamC*, Δ*bamB* TraDIS libraries, listed from innermost tracks outwards, generated using DNAPlotter. The outermost track marks the BW25113 genome in base pairs with the inner grey bar tracks corresponding to sense and antisense CDS.

**Figure S5.**
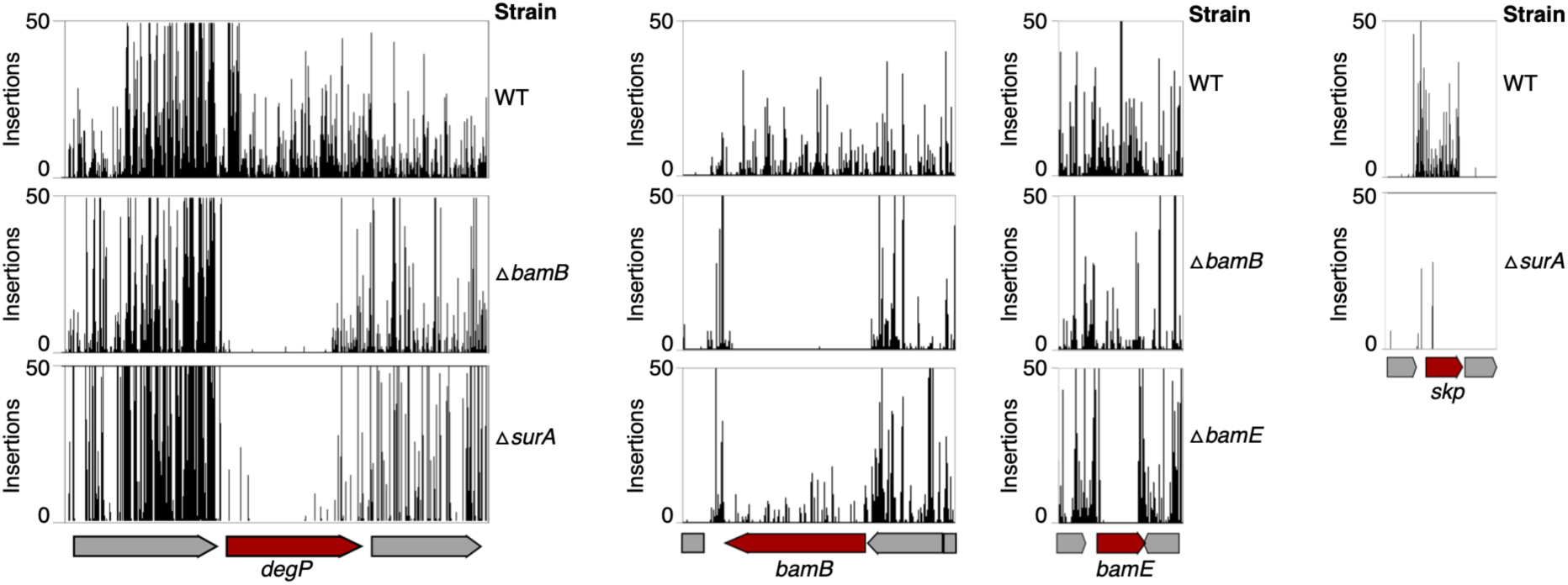
Examples of known synthetic lethal interactions. Transposon insertions in the genes *degP*, *skp*, *bamB* or *bamE* in the WT parent strain, Δ*bamB*, or Δ*surA* mutant TraDIS libraries. Synthetic-lethal genes are represented as red arrows, non-essential genes are represented by grey arrows. Transposon cut-off is set to 50.

**Figure S6.**
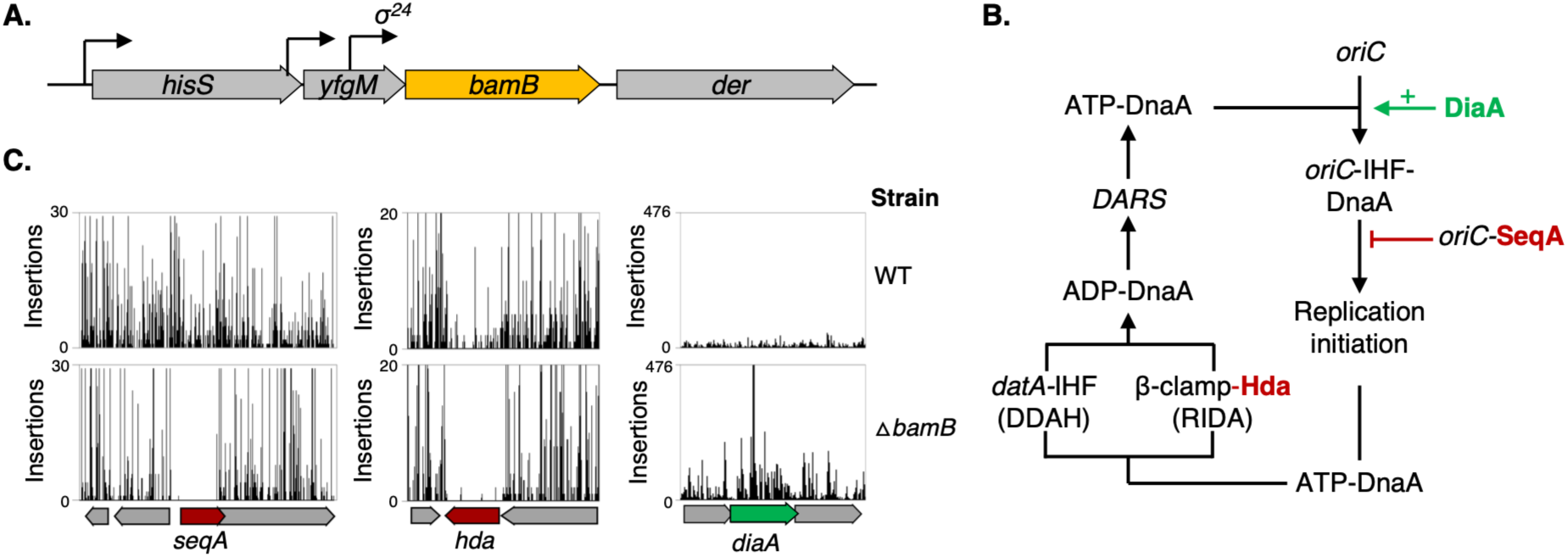
TraDIS identifies genes involved in DNA replication control as essential in the Δ*bamB* strain. **A.** Schematic showing gene organisation and local genetic context at the *bamB* locus. **B.** Schematic representing DNA replication control in *E. coli*. DiaA stimulates initiation at the origin of replication, *oriC*, by DnaA. SeqA prevents premature reinitiation of replication. Hda acts together with the β-sliding clamp to hydrolyse ATP-bound DnaA by regulatory inactivation of DnaA (RIDA). This can also occur through *datA*-dependent DnaA-ATP hydrolysis (DDAH). **C.** Transposon insertions in the genes *hda*, *seqA* and *diaA* in the parent or Δ*bamB* mutant TraDIS libraries. Synthetic-lethal genes are represented as red arrows and genes that increase fitness are in green.

**Figure S7.**
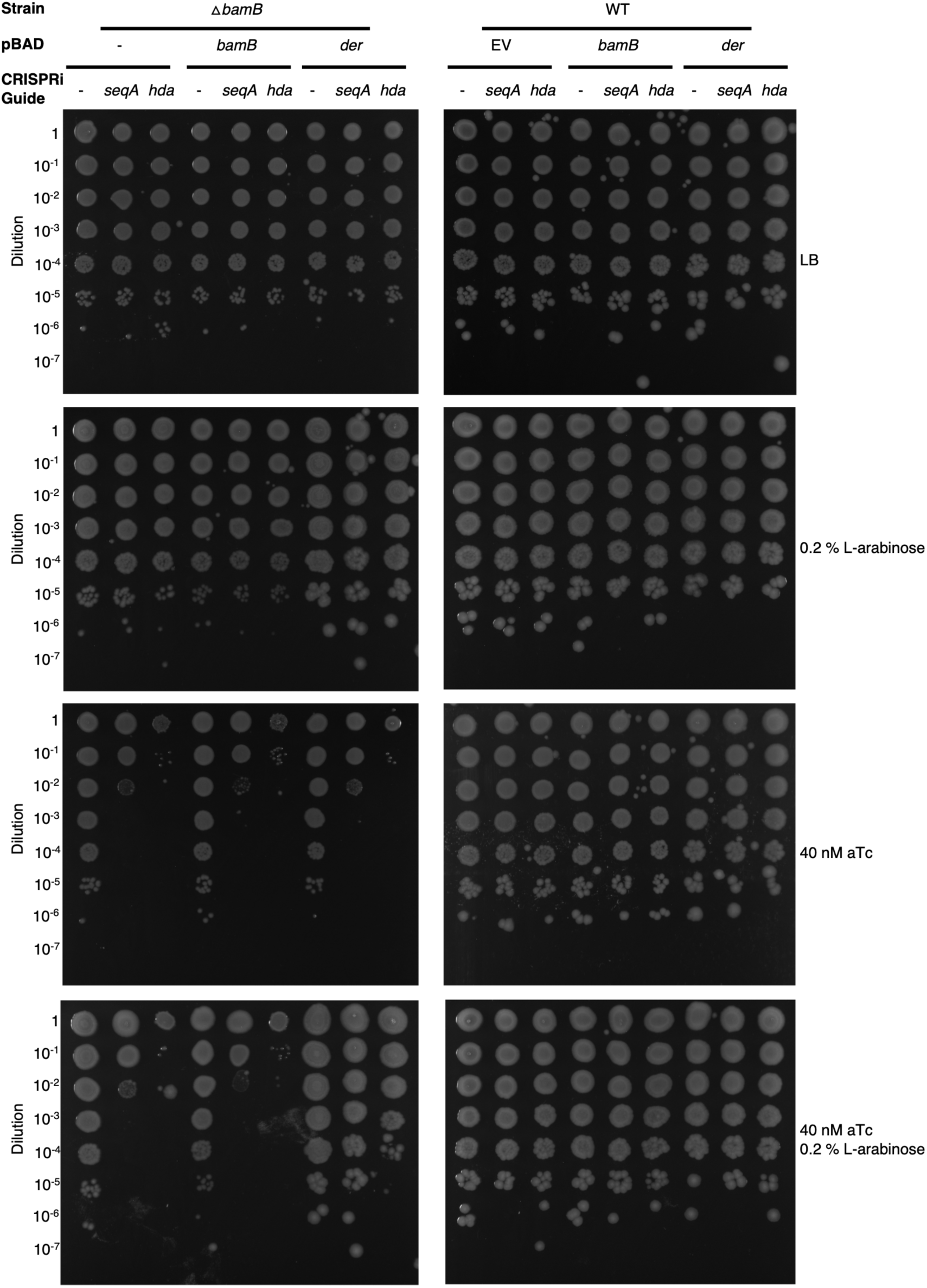
Synthetic-lethality of *seqA* and *hda* in the *bamB* strain is due to effects on expression of *der*. Efficiency of plating assay showing survival of the parent dCas9 expressing strain LC-E18 (WT) or Δ*bamB* mutant derivatives carrying the pSGRNA plasmid, which encodes CRISPRi guide RNA targeting either *seqA* or *hda*. Cells also carry L-arabinose inducible pBAD plasmids encoding *bamB*, *der* or the empty vector control (-). Cells were grown on LB with/without 40 nM anhydro-tetracycline (aTc) to induce guide RNA expression, and with/without 0.2 % L-arabinose to induce expression of genes encoded on the pBAD plasmids.

**Figure S8.**
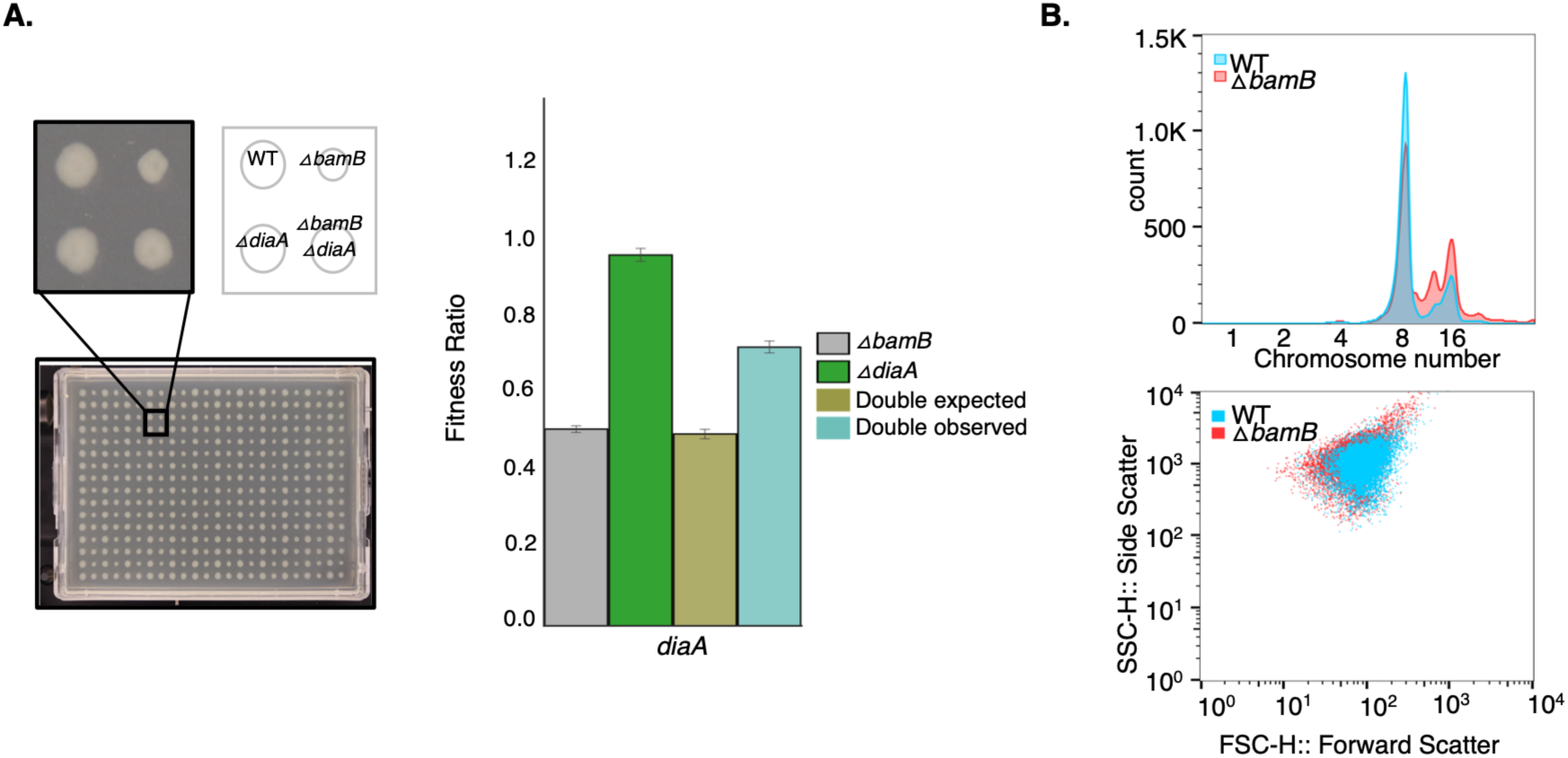
Disruption of *der* expression can be alleviated by *diaA* disruption and affects DNA replication control. **A**. The *E. coli* BW25113 parent strain, single, or double Δ*bamB* and Δ*diaA* mutants were spotted on LB agar in 384-well format, in triplicate, and grown overnight at 37°C before being imaged. Fitness was calculated based on colony size and fitness ratios generated relative to the WT parent. **B.** Flow cytometry of *E. coli* LC-E18 cells grown in LB followed by replication run-out assay. Forward scatter and side scatter are plotted with WT cell data indicated in blue and Δ*bamB* data points in red. Fluorescence is plotted and represents chromosomal content for each cell with chromosome numbers for each peak marked.

**Figure S9.**
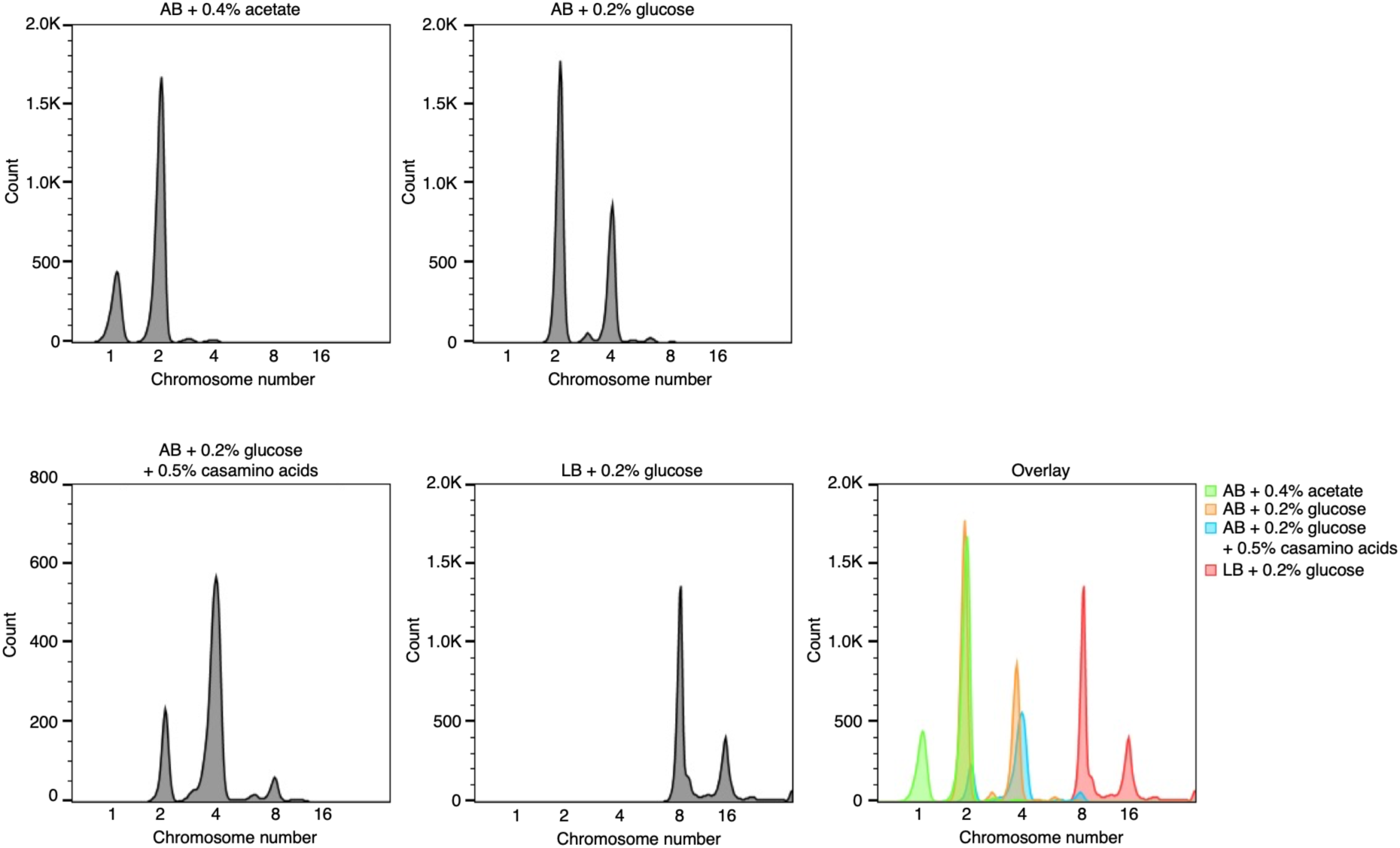
Replication run-out assay standards. Flow cytometry of *E. coli* MG1655 cells grown in media supporting different growth rates followed by replication run-out assay. Cells were grown in AB minimal medium supplemented with either 0.4% acetate, 0.2% glucose, or 0.2% glucose + 0.5% casamino acids, or LB supplemented 0.2% glucose at 37°C with aeration before being treated with rifampicin, cephalexin and stained with Sytox Green. Fluorescence is plotted and represents chromosomal content for each cell with chromosome numbers for each peak marked. An overlay of the separate data panels is presented.

**Figure S10.**
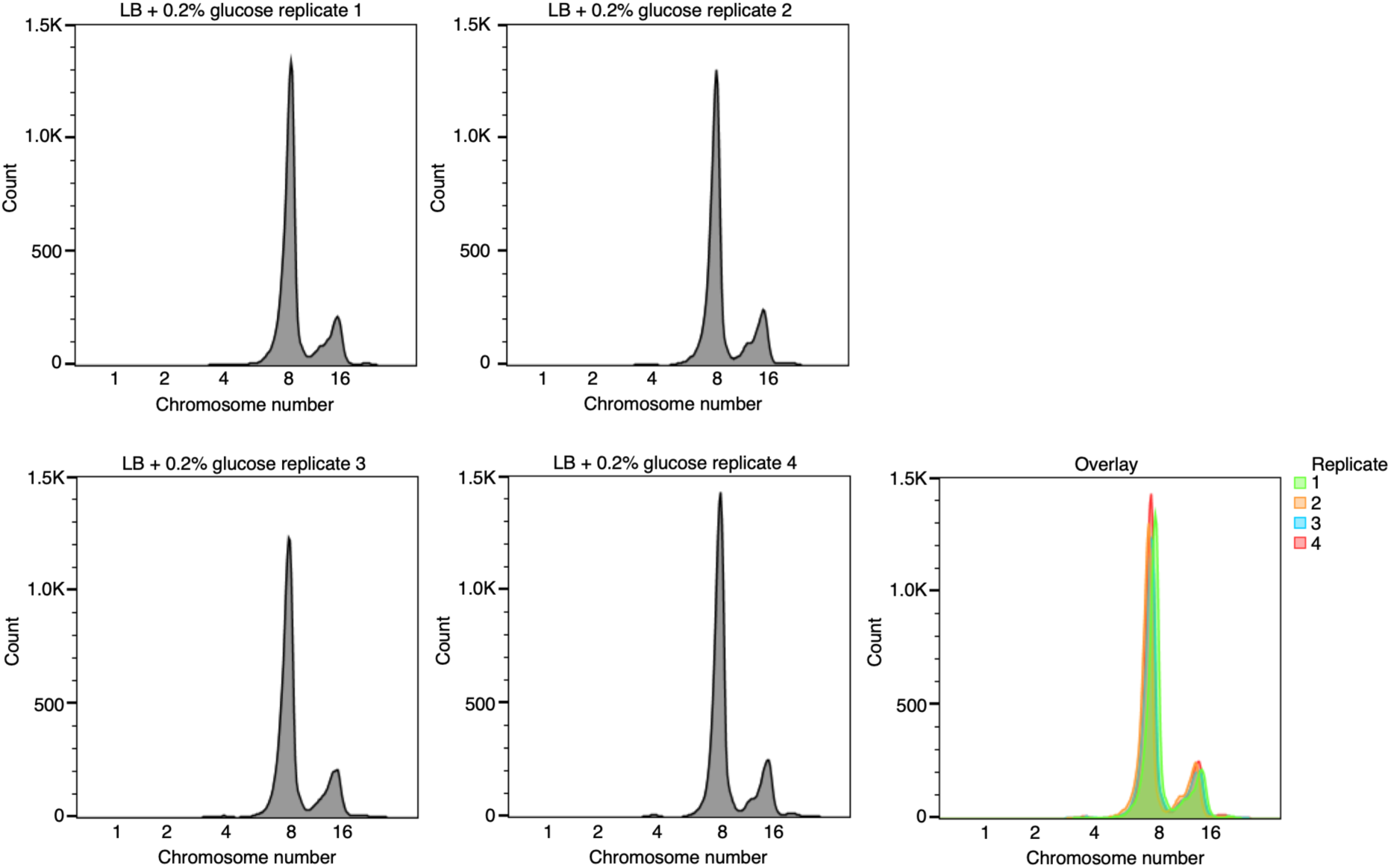
Reproducibility of replication run-out assays for the parent strain. Flow cytometry of *E. coli* LCE-18 cells grown in LB supplemented with 0.2% glucose at 37°C with aeration before being treated with rifampicin, cephalexin and stained with Sytox Green. Fluorescence is plotted and represents chromosomal content for each cell with chromosome numbers for each peak marked. Each experiment was repeated on 4 separate occasions and an overlay of the separate data panels is presented.

**Figure S11.**
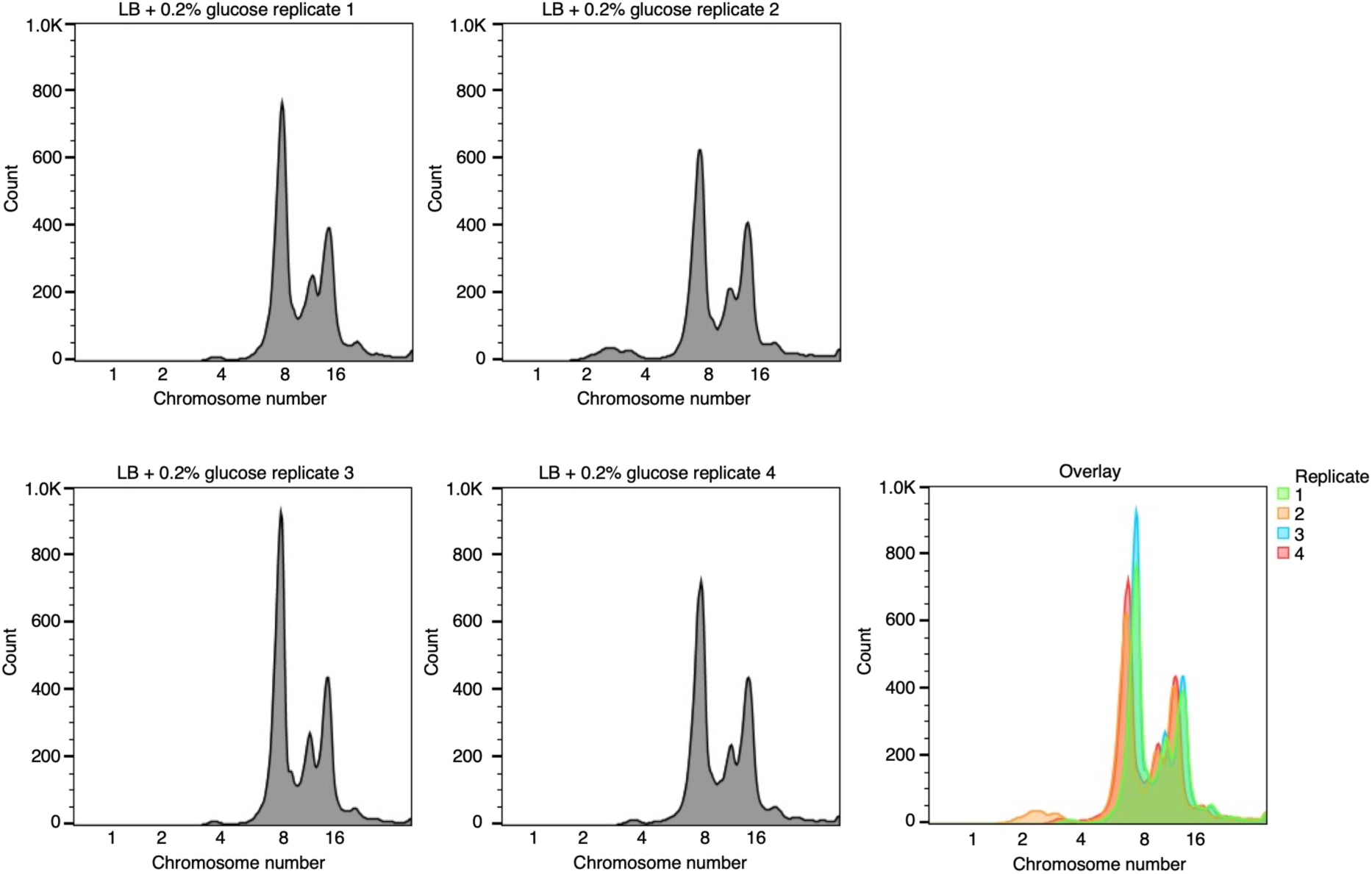
Reproducibility of replication run-out assays for the Δ*bamB* strain. Flow cytometry of *E. coli* LCE-18 Δ*bamB* cells grown in LB supplemented with 0.2% glucose at 37°C with aeration before being treated with rifampicin, cephalexin and stained with Sytox Green. Fluorescence is plotted and represents chromosomal content for each cell with chromosome numbers for each peak marked. Each experiment was repeated on 4 separate occasions and an overlay of the separate data panels is presented.

**Figure S12.**
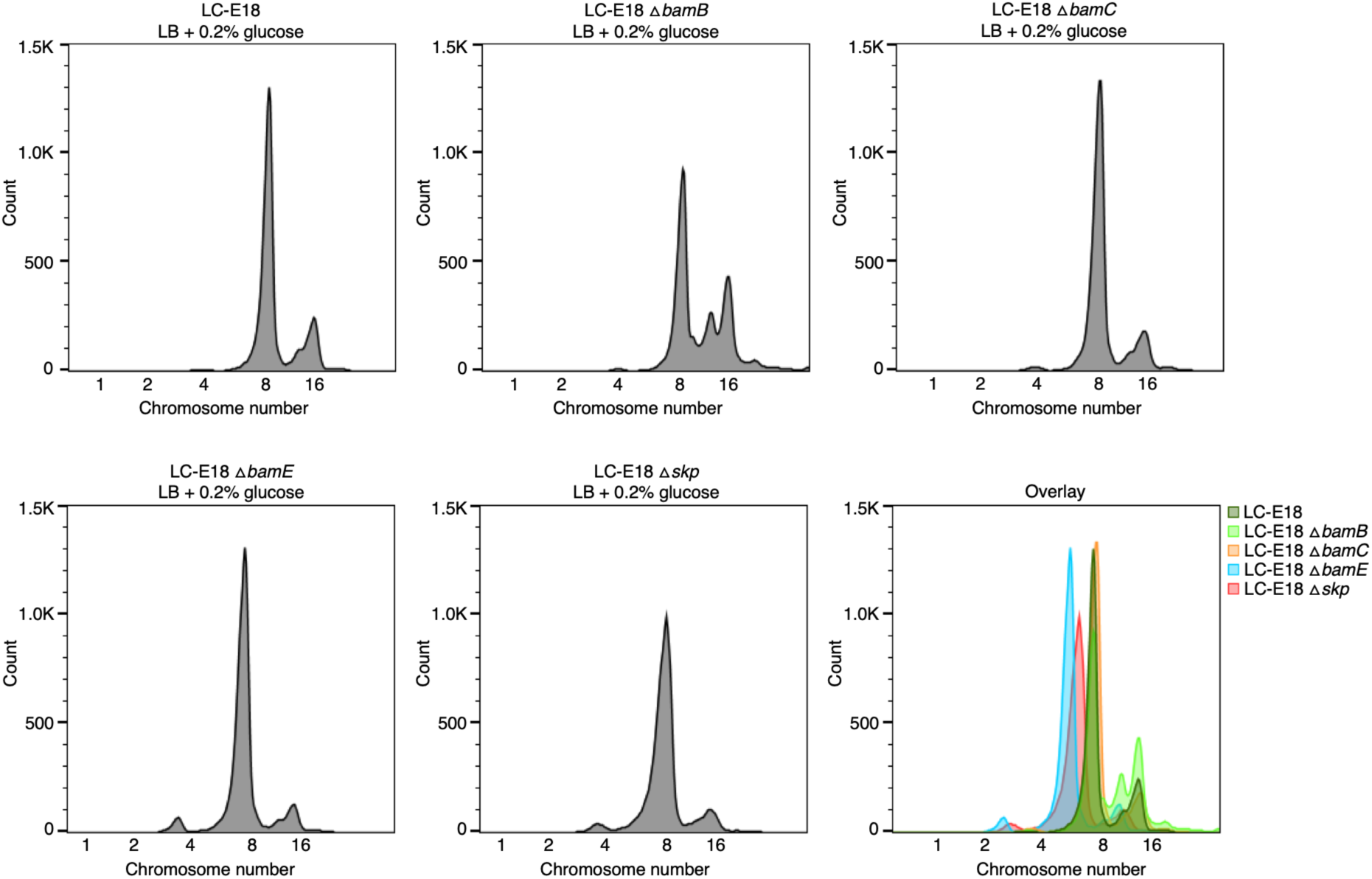
Replication run-out assays for the parent, Δ*bamB,* Δ*bamC,* Δ*bamE* and Δ*skp* strains. Flow cytometry of *E. coli* LCE-18 WT parent, Δ*bamB,* Δ*bamC,* Δ*bamE* or Δ*skp* cells grown in LB supplemented with 0.2% glucose at 37°C with aeration before being treated with rifampicin, cephalexin and stained with Sytox Green. Fluorescence is plotted and represents chromosomal content for each cell with chromosome numbers for each peak marked. An overlay of the separate data panels is presented.

**Figure S13.**
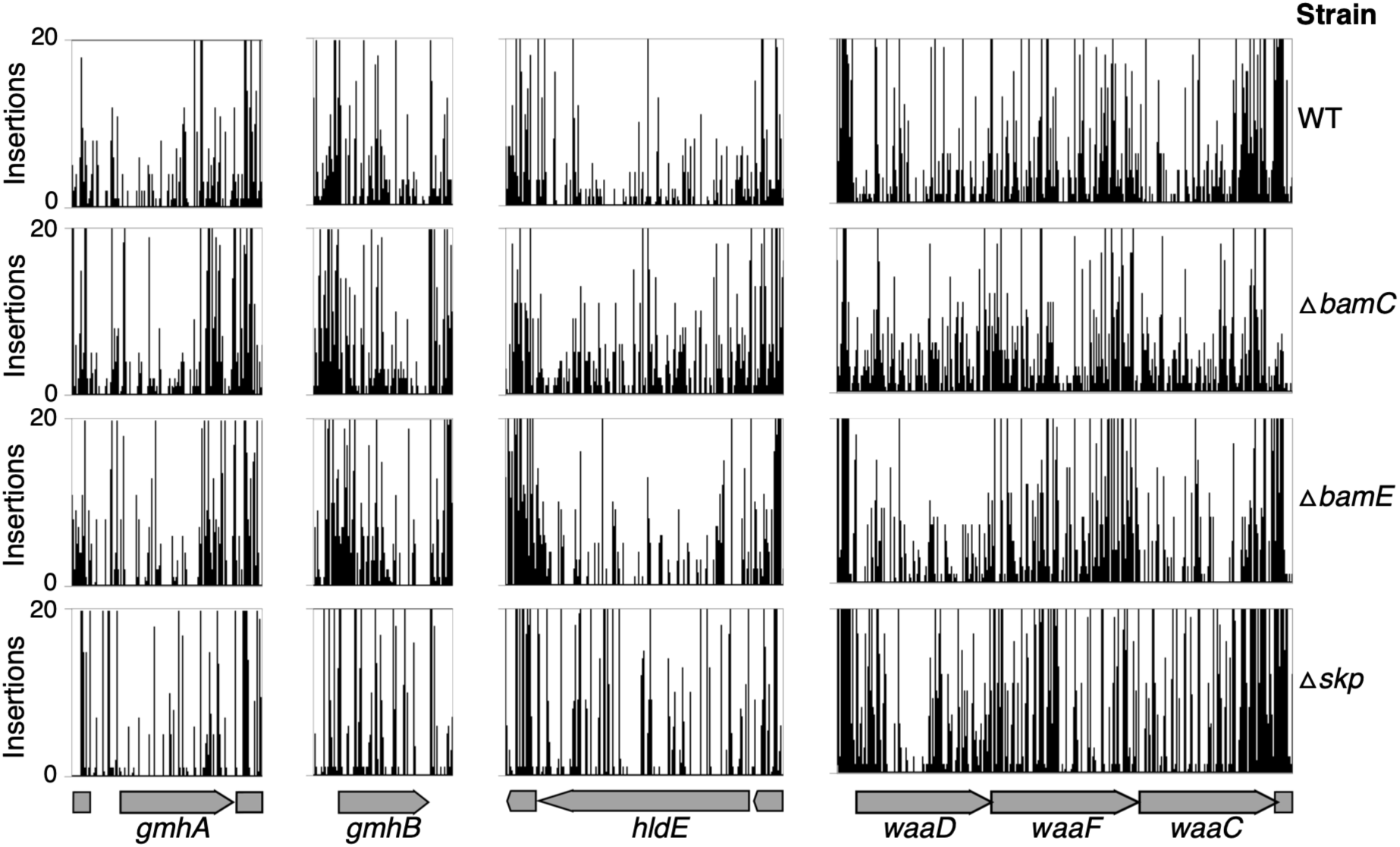
Transposon insertions in genes required for heptose biosynthesis in the Δ*bamC,* Δ*bamE* and Δ*skp* TraDIS libraries. Transposon insertions in the genes *gmhA*, *gmhB*, *hldE, waaD, waaC,* and *waaF* in the parent, Δ*bamC,* Δ*bamE* and Δ*skp* TraDIS libraries. Transposon cut-off is set to 20. Essential genes are represented as red arrows and non-essential genes are represented by grey arrows.

**Figure S14.**
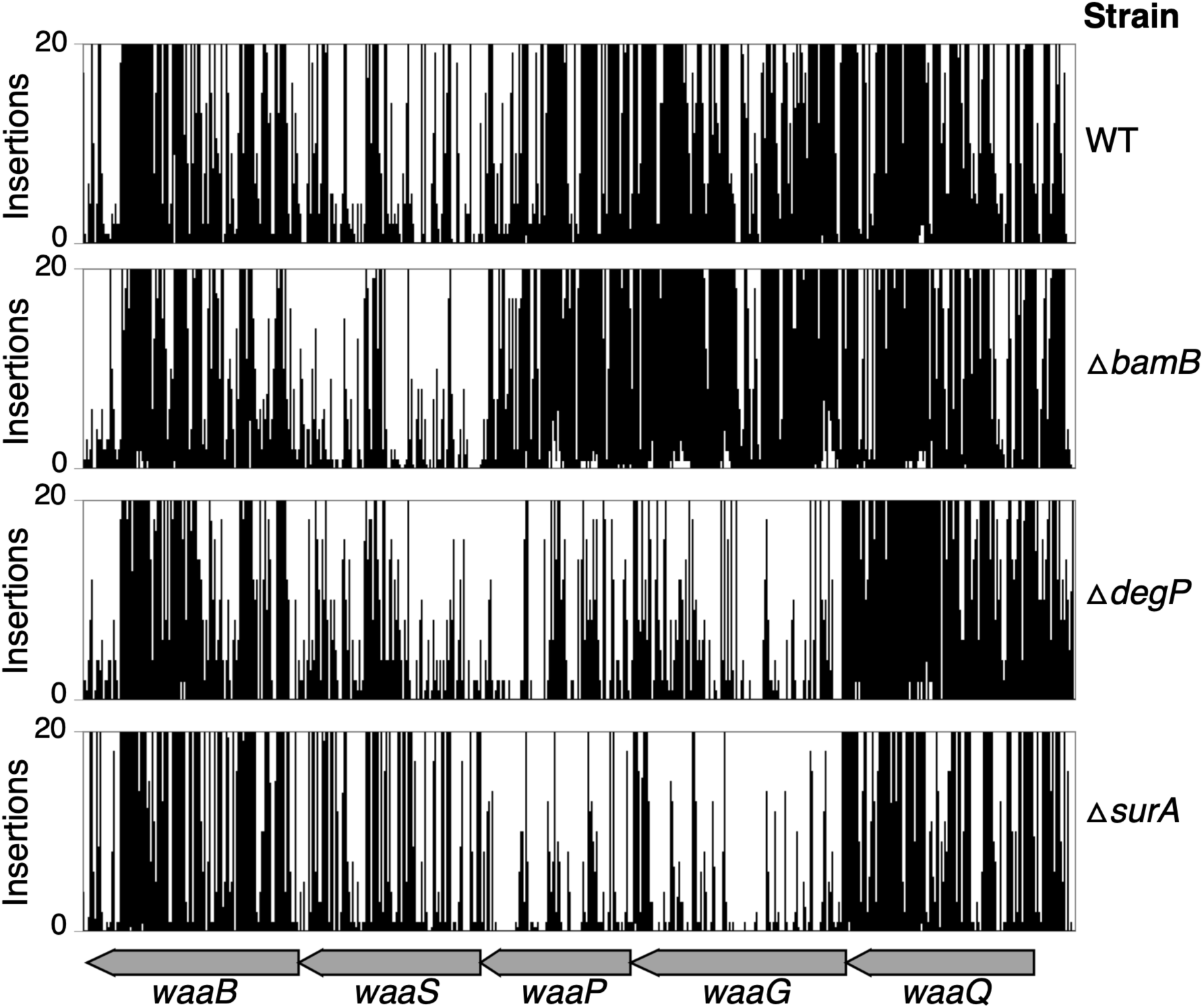
Transposon insertions in the *waaG* and *waaP* genes in the WT, Δ*bamB,* Δ*degP* and Δ*surA* TraDIS libraries. Transposon insertions in the genes *waaB, waaS, waaP, waaG* and *waaQ* in the parent, Δ*bamB,* Δ*degP* and Δ*surA* TraDIS libraries. Transposon cut-off is set to 20. Genes are represented by grey arrows.

## Materials and methods

### Bacterial strains and culture conditions

For TraDIS experiments and phenotypic profiling, the parent strain was *E. coli* K-12 BW25113. The *E. coli* Δ*bamB*, Δ*bamC*, Δ*bamE*, Δ*surA*, Δ*skp* and Δ*degP* mutants were generated by transferring the relevant allele from the Keio library [91] into the clean parent strain by P1 transduction [92]. The *kan^R^* cassette was then removed by using the pCP20 plasmid [93] to leave a small in-frame 34 amino acid peptide consisting of residues from the FRT scar and the first amino acid and last seven of the target gene. These strains were then used for construction of transposon libraries and phenotypic profiling. The same approach was used for construction of the *E. coli* Δ*wecA*::*aph*, Δ*wecD*::*aph*, Δ*wecF*::*aph*, Δ*ompT::aph,* Δ*dapF*::*aph* and Δ*diaA::aph* mutants. The *diaA::aph* allele was also transferred into the Δ*bamB* derivative of *E. coli* BW25113 to generate the Δ*bamB*Δ*diaA::aph* double mutant. The *dapF::aph* allele was also transferred from the Keio library into Δ*bamB*, Δ*bamC*, Δ*bamE*, Δ*surA*, Δ*skp* and Δ*degP* mutants in the presence of 1 mM *meso*-diaminopimelate in order to generate double mutants. The Δ*rcsF*Δ*lpp*Δ*pgsA*Δ*surA* mutant was generated by transfer of alleles from the Keio library and subsequent removal of the *kan^R^* cassette by use of the pCP20 plasmid before incorporation of the next allele by P1 transduction. The *E. coli* Δ*gmhA*::*aph,* Δ*gmhB*::*aph,* Δ*hldE*::*aph,* Δ*waaD*::*aph,* Δ*waaC*::*aph,* Δ*waaF::aph, ΔwaaP::aph,* Δ*waaG::aph,* Δ*waaY::aph* mutants were created by using λ-Red recombination, as previously described for single-step gene inactivation [94]. All mutants were confirmed by polymerase chain reaction and strains were routinely cultured on LB agar and LB broth. The dCas9 expressing strain *E. coli* LC-E18 and the pSGRNA plasmid was a gift from David Bikard (Addgene plasmid # 115924) and has been described previously [42]. The *E. coli* LC-E18 Δ*bamB* derivative was constructed as described for the BW25113 Δ*bamB* derivative with the *kan^R^* cassette removed in order to allow selection of the pSGRNA plasmid in this background. All strains used in this study are listed in **Table S3**. Bacterial cultures were grown at 37°C unless otherwise stated. Where stated, the medium was supplemented with kanamycin (50 μg/ml), carbenicillin (100 μg/ml), *meso*-DAP (1 mM), anhydro-tetracycline (40 nM), or L-arabinose (0.2 %). For micro-dilution survival assays, bacteria were grown in 5 ml LB medium at 37°C with aeration for ∼16 hours. Cultures were normalised by optical density to OD_600_ = 1.00, 10-fold serially diluted in LB, and 2 μl of each dilution was inoculated onto LB agar plates.

### Phenotypic screening

The *E. coli* K-12 BW25113 parent strain, Δ*bamB*, Δ*bamC*, Δ*bamE*, Δ*surA*, Δ*skp* and Δ*degP* mutants were screened against diverse stress conditions to phenotypically profile the mutants. Strains were arrayed in 384-well format and inoculated on 2% agar LB plates using a BM3-BC robot (S&P Robotic Inc.). In addition, the mutants and the wild type were subsequently inoculated on LB agar plates containing different stress conditions. The inoculated plates were incubated for 12 to 14 hours at 37^°^C before being imaged under controlled lighting with an 18-megapixel Canon rebelT3i (Canon) camera on the BM3-BC robot (S&P Robotic Inc.). Images of the plates were analysed using the software IRIS, which measured the size, opacity and circularity of each colony [27]. A total of 4 replica plates were generated for each stress condition. Fitness of the mutants was then scored and analysed using the ChemGAPP Small software [28]. Mean colony size for the mutant in each condition was compared to the mean colony size of that mutant in the LB agar condition, which was normalised to a fitness score of 1 [28]. Fitness scores below 1 represent decreased fitness, as a function of colony size compared to growth on LB agar, and scores above 1 indicate increased fitness in that condition. Each of the plates contained 56 replicates for the parent BW25113 strain (WT), Δ*bamB*, Δ*bamC*, Δ*bamE* and Δ*degP* strains with 52 replicates for Δ*surA* and Δ*skp* for plate space requirements. There were 4 replicate plates for each condition, all of which were treated as individual replicates and compared to their respective wild type parent replicates on each plate before the average was taken. The probability that the two means are equal across replicates was obtained by a one-way ANOVA. Correlation of fitness ratios for each strain was assessed by calculating the Pearson correlation coefficient for averaged fitness scores across replicates using Python before being plotted as a heatmap.

### TraDIS library construction

The *E. coli* K-12 BW25113 parent strain, Δ*bamB*, Δ*bamC*, Δ*bamE*, Δ*surA*, Δ*skp* and Δ*degP* mutants were transformed with the EZ-Tn5™ <KAN-2> Tnp Transposome (Cambio) as previously described [32]. Approximately 1 million mutant colonies were pooled, thoroughly mixed and stored in LB supplemented with 15% glycerol at -80°C. DNA was extracted from at least two samples of each transposon library to generate independent sampling replicates for library generation.

### Sequencing of TraDIS libraries

Cells from the pooled mutant library were harvested and genomic DNA was extracted for library preparation and sequencing as previously described [32]. Samples were sequenced using an Illumina MiSeq with a 150 cycle v3 cartridge. Raw data is available at the ENA under accession PRJEB121496.

### TraDIS data analysis

Raw data were processed using a series of custom scripts as previously described [32]. The data were trimmed using Fastx barcode splitter and trimmer tools (Pearson *et al.,* 1997) and filtered based on inline indexes. The accuracy of the transposon sequences were checked in two steps: the first 22 bases, allowing for three nucleotide base mismatches and the last 10 bases of the transposon, allowing for up to one mismatch. Using Trimmomatic, sequences with less than 20 bases in length were removed (Bolger *et al.,* 2014). TraDIS data were then analysed using BioTraDIS (https://sanger-pathogens.github.io/Bio-Tradis/) [34] and aligned to the *E. coli* BW25113 reference genome CP009273.1, available from NCBI (Tatusova *et al.,* 2014). We used SMALT (https://www.sanger.ac.uk/tool/smalt/) as an aligner with zero value for mismatch threshold. We also set the parameter smalt_r, determining how to treat multi-mapping reads, to zero. This avoided repetitive elements to count as essential. We ran the tradis_essentiality.R script within the BioTraDIS package independently on each library: the six mutant backgrounds (Δ*bamB,* Δ*bamC,* Δ*bamE,*Δ*surA,*Δ*skp,*Δ*degP*) and two *E. coli* BW25113 WT reference sets (an “internal” WT replicate sequenced as part of this study, and an “external” WT dataset from a previous study[95]). This classifies each gene as essential, ambiguous, or non-essential for each library based on the bimodal distribution of insertion indices [30, 34]. Synthetic-lethal gene lists were then built by comparing essentiality classifications between each mutant and the WT sets, which were then flagged as shared/not shared with the internal or external WT essential gene lists in Supplementary Table 1. Therefore, a gene was treated as synthetic-lethal in a given mutant when it was called essential in that mutant but not shared with the WT essentiality call.

### Functional enrichment analysis

KEGG pathway enrichment analysis was performed on the synthetic-lethal gene sets for each mutant background using the enrichKEGG function from the clusterProfiler R package [96], with the whole E. coli K-12 BW25113 genome used as the background gene set. Gene ratio is defined as the proportion of genes within a given synthetic-lethal gene set that are annotated to a specific KEGG pathway. Enrichment significance was assessed using a hypergeometric test comparing pathway representation within each query gene set to the whole-genome background.

### Membrane fluidity assay

Membrane fluidity was measured using the membrane fluidity kit (Abcam: ab189819), as previously described, but with minor modifications [97]. Bacterial strains were grown to mid-exponential phase (OD_600_ = ∼0.4-0.6) in LB medium. Cells were harvested by centrifugation, washed with phosphate buffered saline (PBS) and incubated with labelling mix (10 μM pyrenedecanoic acid (PDA), 0.08% pluronic F-127, in PBS) in the dark for 20 min at 25°C with rocking. Cells were then washed twice in PBS and re-suspended in PBS prior to measuring fluorescence (excitation = 350 nm, emission = either 400 nm or 470 nm). Membrane fluidity was estimated by measuring the ratio of excimer (470 nm) to monomer (400 nm) fluorescence. The emission spectra were compared to unlabelled cells to confirm membrane incorporation and each experiment contained triplicate technical repeats and was then repeated three times. Membrane fluidity of the mutants of interest were expressed relative to the parent *E. coli* strain. Experiments were performed in technical triplicate and were repeated three times. Two sample t-tests were used to assess statistical significance of differences from the WT strain. Raw data are available in supplementary data file 4.

### OmpT *in vivo* fluorescence assay

The OmpT assay for monitoring Bam activity was performed as described previously with minor modifications [10, 16, 47]. Bacterial strains were grown to mid-exponential phase (OD_600_ = ∼0.4-0.6) in LB medium and were normalised to OD_600_ of 0.2 in growth media. The cell suspension (5 μl) was then added to 95 μl of 25 μM fluorogenic peptide, Abz-ARRAY(NO_2_)-NH_2,_ diluted in PBS. Fluorescence emission was immediately measured (excitation = 325 nm, emission = 430 nm) over a period of 5 h, with readings every 20 s. OmpT activity was expressed relative to the parent strain. Experiments were performed in technical triplicate and were repeated three times. Two sample t-tests were used to assess statistical significance of differences from the WT strain. Raw data are available in supplementary data file 4.

### Phospholipid extraction and thin layer chromatography

Phospholipids were extracted using an adapted Bligh-Dyer method [98]. Bacterial cultures were grown overnight (∼16 hours) at 37°C with shaking and cells were harvested by centrifugation. The bacterial cell pellets were resuspended and normalised to OD_600_ = 3 and mixed with methanol and chloroform (2:1). The cell suspension was incubated at 50°C for 30 min before additional chloroform and water was added to the samples to produce a final ratio 2:2:1.8 of methanol/chloroform/water. Following centrifugation, the phospholipid-containing phase was extracted and dried under nitrogen before being stored at -20°C. Samples were re-dissolved in chloroform before being separated by thin layer chromatography on silica gel membrane using a solvent system that consisted of chloroform, methanol, acetic acid (65:25:10). The plate was subsequently stained with phosphomolybdic acid (PMA) and warmed until the lipid species were visible.

### Efficiency of plating assay

Efficiency of plating assays were completed as previously described [99]. Cells were grown overnight in LB (supplemented with 1 mM *meso*DAP where indicated) before being normalised to OD_600_ = 1.00 and serially diluted 1:10. Following dilution, 2 μl was spotted on agar plates containing supplements where indicated and incubated at 37°C for ∼16 hours before being imaged.

### Genetic interaction analysis

For genetic interaction analysis, overnight cultures were diluted 1/100 and then grown to OD_600_ = 0.8-1. Cultures were then spread on LB agar plates using sterile glass beads to create source plates, before being incubated for 12 to 14 hours at 37^°^C. Each strain was arrayed on LB agar plates, from the source plates, to form 384 colonies of 96 replicates for each strain using a BM3-BC robot (S&P Robotic Inc.). Plates were incubated at 37^°^C for 12 to 14 hours. The plate was then imaged under controlled lighting, with an 18-megapixel Canon rebelT3i (Canon) camera on the BM3-BC robot (S&P Robotic Inc.). Images were then analysed using the software called IRIS to measure the size, opacity and circularity of each colony on the plates [27]. Fitness of the mutants was then analysed and compared using the software ChemGAPP GI [28].

### Flow cytometry analysis

Chromosome number measurements were performed as described previously, but with several modifications [43]. Briefly, cells were grown with aeration at 37°C until OD_600_=0.15 in LB medium supplemented with 0.2% glucose. Samples were collected, treated with 150 μg/ml rifampicin, and 10 μg/ml cephalexin and incubated for 4 h at 37°C with mixing. Incubation with antibiotics results in cells containing an integral number of chromosomes, corresponding to the number of replication origins at the time of drug treatment. Subsequently, cells were harvested, washed with TBS (20 mM Tris-HCl pH 7.5, 130 mM NaCl) and fixed with cold 70% ethanol overnight. Additional samples were collected at OD_600_=0.15 without antibiotic treatment and fixed as above.

Prior to flow cytometry analysis, cells were resuspended in 50 mM sodium citrate followed by RNA digestion with RNase A for 4 h. Chromosomal DNA was stained with 2 mM Sytox Green (Invitrogen) and DNA content per cell was measured with BD FACS Calibur at 488 nm Argon Ion laser. MG1655 (WT) strain grown in AB medium containing one of the following carbon sources: 0.4% sodium acetate, 0.2% glucose, 0.2% glucose + 0.5% casamino acids or in LB medium with 0.2% glucose, treated with antibiotics, fixed and stained as above was used as a standard for each flow cytometry measurement, indicating cells containing 1/2, 2/4, 4/8 or 8/16 chromosomes, respectively. Flow cytometry data was analyzed using FlowJo ver. 10.8.0.

### Conservation analysis

Reference genomes were downloaded for each species from NCBI in fasta format and annotated using Prokka [100]. The 16s rRNA sequence for each species was then extracted from Prokka ffn files in fasta format. A Kmer based neighbour joining tree was produced via extracting a one-step sliding window of 8 base kmers from each 16s rRNA sequence. Following this, the number of unique kmers was counted for each species in order to produce a Kmer profile. The Jaccard similarity between each species Kmer profiles was then used to produce a distance matrix. This distance matrix was input into the R module ape::nj to produce the neighbour joining tree. The neighbour joining tree was reformatted as a nexus file and visualised within Figtree (http://tree.bio.ed.ac.uk/software/figtree/), where the tree was rooted at the midpoint and branches were transformed to become proportional.

To count the instances of the proteins BamA, BamB, BamC, BamD, BamE, DegP, SurA, and Skp, hmmsearch was used to extract proteins with the relevant domains from the Prokka annotated faa files. The Pfam domains used were PF01103, PF13360, PF06804, PF13525, PF04355, PF09312 and PF03938 for BamA, BamB, BamC, BamD, BamE, SurA, and Skp, respectively. For DegP, both PF00595 and PF13180 were used and the intersect taken. An e-value of 0.001 was used as a threshold for significant hits by hmmsearch. For BamA, hits annotated as the query protein were inputted into hmmscan and hits with at least one POTRA domain and an Omp85 domain were counted as true BamA hits. Hits containing a Tam-POTRA domain were excluded. Where a hit had only an Omp85 gene, the adjacent gene was checked for the POTRA domains. For BamB, any hmmsearch hits annotated as BamB or hypothetical proteins by Prokka were input into SignalP 6.0 [71] and hmmscan. Proteins assigned a LIPO (Sec/SPII) signal peptide within SignalP 6.0, with only one PQQ_2 domain within hmmscan were selected as true BamB hits. For BamC, all hits from hmmsearch were inputted into hmmscan to confirm the presence of the Lipoprotein-18 domain. Hits were also inputted into Phmmer and proteins with significant homology to BamC were counted as true hits. For BamD, hypothetical proteins and BamD hits were inputted into hmmscan and Phmmer and those with homology to BamD and a non-outcompeted YfiO domain counted as true hits. For BamE, hypothetical proteins and BamE hits were inputted into hmmscan and those with a singular SmpA_OmlA domain were counted as true hits. For DegP hypothetical proteins and DegP hits were inputted into hmmscan. If a hit had a PDZ type domain, Peptidase_M50 domain or Trypsin_2 domain then it was visualised on Pfam and manually assigned as DegP based on structure. If a DegP hit had any other type of domain, it was discounted. For SurA, all hits were run through hmmscan and Phmmer. Those with a SurA_N domain, Rotamase domain and a Rotamase_3 domain and homologous to SurA within Phmmer were counted. For Skp, all hits were run through Phmmer and only hits with only an OmpH domain and with homology to Skp counted as true hits. Finally, the neighbour-joining tree was edited in iTOL [73] to produce the heatmap of gene counts.

